# Valine overproduction with metabolically engineered *Methanothermobacter marburgensis*

**DOI:** 10.64898/2025.12.09.693116

**Authors:** François Unger, Sophie Marichez, Maximilian Klein, Ross T. Fennessy, Justin Smith, Thomas Stehrer-Polašek, Simon K.-M. R. Rittmann, Christian Fink

**Author notes:** **Correspondence:** Christian Fink,; Simon K.-M. R. Rittmann.

## Abstract

In times of a climate crisis caused by extreme emissions of green-house gases on a global scale, mitigation solutions need to be found. One solution is the system of carbon capture and utilization (CCU), where C1 gases, such as carbon monoxide (CO), carbon dioxide (CO_2_), or methane, are either redirected from industrial off-gas streams or directly air-captured. A biotechnological process for CCU is the use of *Methanothermobacter marburgensis* for CO_2_ fixation and production of value-added compounds. In this study, we focused on valine production, an amino acid important for human or feed stock nutrition. We demonstrated overproduction of valine from CO_2_ in *M. marburgensis* with temperature-induced promoters. Here, we reached a 12.9-fold increase in valine production between the OFF- and ON states of the inducible promoter with a maximal specific production rate of 14.17 mg gCDW^-1^ h^-1^ of valine in closed batch experiments. In the second approach for valine production, we overexpressed acetolactate synthase genes with resistance to allosteric valine inhibition from *Methanothermobacter thermautotrophicus* recombinant in *M. marburgensis*. We identified a strong reduction in allosteric inhibition towards valine. This resulted in specific valine productivity of up to 40 mg gCDW^-1^ h^-1^ and states the highest specific productivity on an individual amino acid in methanogens. With those findings, we expanded the toolbox for genetic modification of *M. marburgensis* by a thermo-inducible promoter system and applied protein engineering for enhanced production of value-added compounds to *M. marburgensis*. This proof of concept shows the feasibility of archaeal cell factories generation via genetic engineering for industrial production of value-added compounds with thermophilic methanogens.

## Introduction

The use of carbon dioxide (CO_2_) for production of commodity chemicals usually made from fossil resources has the potential to slow down anthropogenic climate change by removal of CO_2_ from the atmosphere and reduction of fossil resources (Ramirez-Corredores 2024). Reduction of CO_2_ to commodity chemicals has been demonstrated with biological catalysts (Liu et al. 2020). In this field, acetogenic bacteria (acetogens) and methanogenic archaea (methanogens) are in the spotlight due to their ability to efficiently fixate CO_2_ and naturally produce valuable compounds, such as acetate, ethanol, or renewable energy in form of methane (Liew et al. 2016; Jin et al. 2020; Martin et al. 2013; Rittmann et al. 2018; Pfeifer et al. 2021). Processes to harness acetogens and methanogens as cell factories for those compounds from CO_2_ are already developed to (pre-) commercial scales in companies, such as LanzaTech Inc., Electrochaea GmbH, or Krajete GmbH (Angenent et al. 2022; Liew et al. 2022). Acetate, ethanol, and methane are terminal catabolic products of the energy metabolism of acetogens or methanogens and therefore naturally produced in high quantities (Liew et al. 2022). Microbial cell factories for more complex compounds derived from anabolic fixated CO_2_, however, are possible but require genetic, metabolic, and enzymatic engineering to result in high production rates and yields (Baumschabl et al. 2022; Contreras et al. 2022; Mühling et al. 2024). First successful proof of concept studies have already been reported for products, such as geraniol and PHB with *Methanococcus maripaludis* (Thevasundaram et al. 2022; Lyu et al. 2016) or isoprene subunits and lactate with engineered *Methanosarcina* sp. (Aldridge et al. 2021; McAnulty et al. 2017). Furthermore, natural secretion of amino acids has been observed in *Methanothermobacter marburgensis* (Taubner et al. 2023; Reischl et al. 2025). This autotrophic, hydrogenotrophic methanogen with an optimal growth temperature of 65 °C is one of the best-studied model organisms in the field of thermophilic archaea (Thauer 2015; Kaster et al. 2011). The fact that *M. marburgensis* shows robust growth and scalability in bioreactor systems makes it a highly suitable candidate for industrial-scale bioprocesses (Rittmann et al. 2018). Together with the recently established tools for genetic engineering, this led to a deepened understanding especially in branched chain amino acids biosynthesis in *M. marburgensis* on a genetic and physiological level (Klein et al. 2025).

Branched chain amino acids (BCAA; valine, leucine, and isoleucine) are essential nutrients for humans and feed stock (Hu et al. 2016). Particularly valine shows a large variety of use cases not only as animal feed or in sports nutrition but also in pharmaceutical, agricultural, and cosmetics applications (Cosmetics Info 2025; Sharma et al. 2024; Cai et al. 2025; Wolfe 2017). While valine can be produced from protein hydrolysates or via chemical synthesis, the main share of approximately 2.65 thousand tons in 2024 that made a market volume of ∼309 million US$ (Chemanalyst 2025; Valuates reports 2025) is produced with glucose-based microbial fermentation using genetically engineered strains of *Escherichia coli* and *Corynebacterium glutamicum* (Gao et al. 2021; Sheremetieva et al. 2024; Wang et al. 2024). Although glucose-based fermentation is still the state-of-the-art for valine production, laboratory-scale production of valine from CO_2_ or CO_2_-derived C1 compounds, such as formate, methanol, or acetate has been demonstrated (Ågren 2024; Wang et al. 2020; Cotton et al. 2020).

Valine biosynthesis starts with deviating carbon flux from the citric acid cycle and pyruvate as the substrates. Four enzymes, including the acetolactate synthase (AHAS, IlvB/N), the ketolacid reductoisomerase (IlvC) and the dihydroxy-acid dehydratase (IlvD) encoded in an operon (IlvBCD), and the branched-chain-amino-acid aminotransferase (IlvE) are involved in its biosynthesis (Oldiges et al. 2014). This biosynthesis pathway is conserved across the three domains of life including methanogenic archaea (Xing and Whitman 1991). While regulatory effects, such as attenuation have a significant impact on expression of the IlvBCD operon, allosteric end-product inhibition of AHAS via valine and leucine appears to have the strongest impact on valine biosynthesis (Elišáková et al. 2005; Gao et al. 2021). On an enzyme level, the release of this allosteric inhibition using enzyme engineering paired with constitutive overexpression of AHAS resulted in largely improved production of valine in *E. coli* and *C. glutamicum* (Eggeling et al. 1987; Elišáková et al. 2005; Morbach et al. 2000; Leyval et al. 2003; Blombach et al. 2009). An exchange of only 1-3 amino acids in the regulatory subunit of AHAS resulted in complete release of feedback inhibition (Elišáková et al. 2005; Park and Lee 2010; Kopecký et al. 1999). This loss of regulation can result in accumulation of intermediates. Therefore, fine-tuning of the expression of downstream genes, such as *ilvC*, *ilvD*, and *ilvE* can be crucial for efficient valine production in industrial scales (Gao et al. 2021).

The overproduction of valine or other compounds in microbial fermentations increases the genetic and metabolic burden on the microorganism. This is often accompanied by reduced growth rates or complete growth inhibition (Mao et al. 2024). Here, inducible systems to split the process into a growth- and production phase can be of great advantage. One way is the use of inducible promoter systems, such as IPTG induced *lac* promoter, arabinose induced *bad* promoter, cumate based systems, and *rhaBAD* promoters induced by rhamnose. (Alagesan et al. 2018). In methanogens native regulatory promoters have also been identified. In *M. maripaludis* native regulatory systems are used in inducible promoters, such as phosphorous or nitrogenated compounds to steer gene expression (Akinyemi et al. 2021; Lie et al. 2005). Furthermore, in *Methanosarcina mazei* protein expression can be induced by methanol (Mondorf et al. 2012). However, the use of native regulatory effects always has an impact on the chassis strain as well. For the use in *Methanosarcina acetivorans*, the non-native tetracycline-based inducible promoter (TetR) system from *E. coli* has been adapted (Guss et al. 2008). This TetR system has also been proven functional in thermophilic autotrophic bacteria with a growth temperature of 65 °C. In this case, the functionality is given even without the need for tetracycline, since the repressor is released naturally from the DNA at a temperature above 60 °C resulting in a temperature-induced promoter system (Mol 2022) .

In this study, we amended this TetR promoter system for usage in *M. marburgensis.* We aimed to both show the capabilities of *M. marburgensis* as a cell factory for inducible valine production from CO_2_ and further development the genetic toolbox for thermophilic methanogens. Therefore, we approached valine overproduction with *M. marburgensis* from two sides, through constitutive gene expression of AHAS with promoters of differing strength but also with enzyme engineering of AHAS towards release of allosteric valine inhibition.

## Materials and methods

All chemicals were obtained from Carl Roth GmbH + Co. KG, Karlsruhe, Germany except stated otherwise.

### Cloning strategy, construct generation, and molecular biology techniques

All cloning strategies were outlined in the SnapGene software (SnapGene software (www.snapgene.com). The primers were designed with a calculated melting temperature of the binding part of 60 °C in the software. Global primer melting temperatures might vary as restriction enzyme recognition sequence might have been added at the 5’ end of the primers. All primers and Sanger sequencing were purchased from Eurofins genomics (Eurofins Genomics Europe Shared Services GmbH, Ebersberg, Germany) and are listed in **table 2**. The synthesized fragment of the codon optimized acetolactate synthase (AHAS) was synthesized by Twist Biosciences (South San Francisco, CA, USA) and delivered in a high copy number ampicillin resistance standard vector (**Supplementary table S1**).

In the first step, we utilized the pArk00005 as a base vector and integrated *Xba*I as additional unique restriction enzyme recognition site between *Avr*II and *Kas*I (Klein et al. 2025). With that, we created two additional modules between *Avr*II and *Kas*I resulting in pArk00007. Afterwards, we exchanged the non-sense DNA between *Kas*I and *Xba*I with the Twist Bioscience synthesized fragments of AHAS0, AHAS2, and AHAS3 resulting in pArk00008, pArk00014, and pArk00015, respectively (Table 1, **Supplementary table S1**). In the next step, we exchanged the *hpt* downstream homologous flank with *tetR* initiated by P_hmtB_ ordered from TWIST Bioscience in an ampicillin resistance containing high copy number vector. Therefore, we digested the P_hmtB__*tetR* fragment and pArk00008 with *Fse*I and *Avr*II and ligated with Instant Sticky-end Ligase Master Mix (New England Biolabs (Ipswich (MA), USA), resulting in pArk00009. The promoters P_hmtB_ _TATA, P_hmtB_ _Trans, P_hmtB_ _TATA_Trans, SIP1 and SIP2 were ordered as DNA fragments with adapters from Twist Biosciences (South San Francisco, CA, USA) (**Supplementary table S1**). The promoter fragments also contained the first 200 bp of the *M. thermautotrophicus* AHAS gene codon-optimized for *M. marburgensis* including the unique *Mfe*I restriction enzyme recognition site. Additionally, we added the unique modular restriction enzyme recognition site *Kas*I upstream of the promoter. With *Mfe*I and *Kas*I restriction enzyme digest of the promoter fragments and pArk00009 according to manufactureŕs manual (New England Biolabs (Ipswich (MA), USA) and ligation with Instant Sticky-end Ligase Master Mix we exchanged the promoters of pArk00009 to generate pArk00010-12 and pArk00016-17 (Table 1). A representative construct is presented in **Supplementary figure S1+S8**. We used *E. coli* NEB5α for standard cloning approaches and storage. Additionally, we generated pArk00013 without promoter upstream of AHAS0. Therefore, we PCR amplified the first 500 bp of AHAS0 from pArk00008 with Q5 high-fidelity polymerase (New England Biolabs (Ipswich (MA), USA) with primers Res_Ark_10-11 (**Table 2**). The resulting fragment and pArk00008 were digested with *Mfe*I and *Kas*I and ligated to yield pArk00013. Competent cells were either used directly from the manufacturer or made chemically competent by suspending them in calcium chloride solution (Sambrook et al. 1989). After confirmation of correct constructs by Sanger sequencing, plasmids were used to chemically transform *E. coli* S17-1 for subsequential conjugation with *M. marburgensis* (**Table 1**).

**Table 1:**
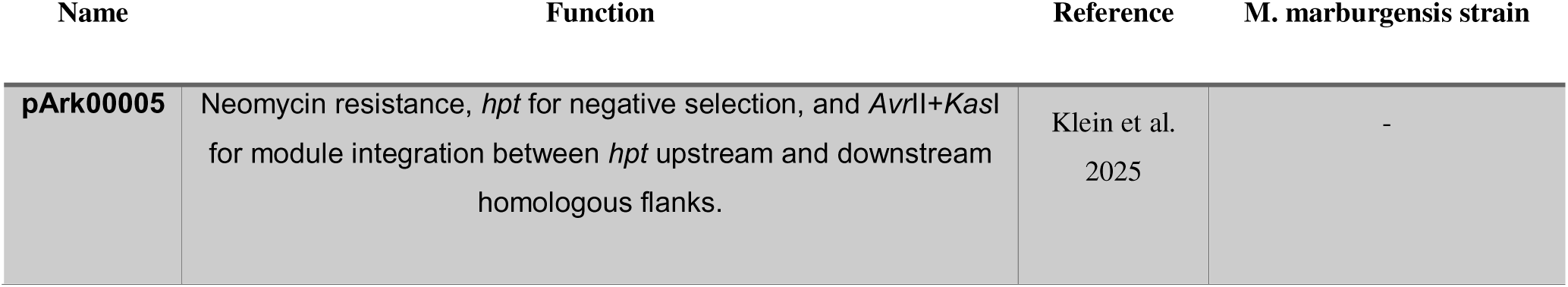
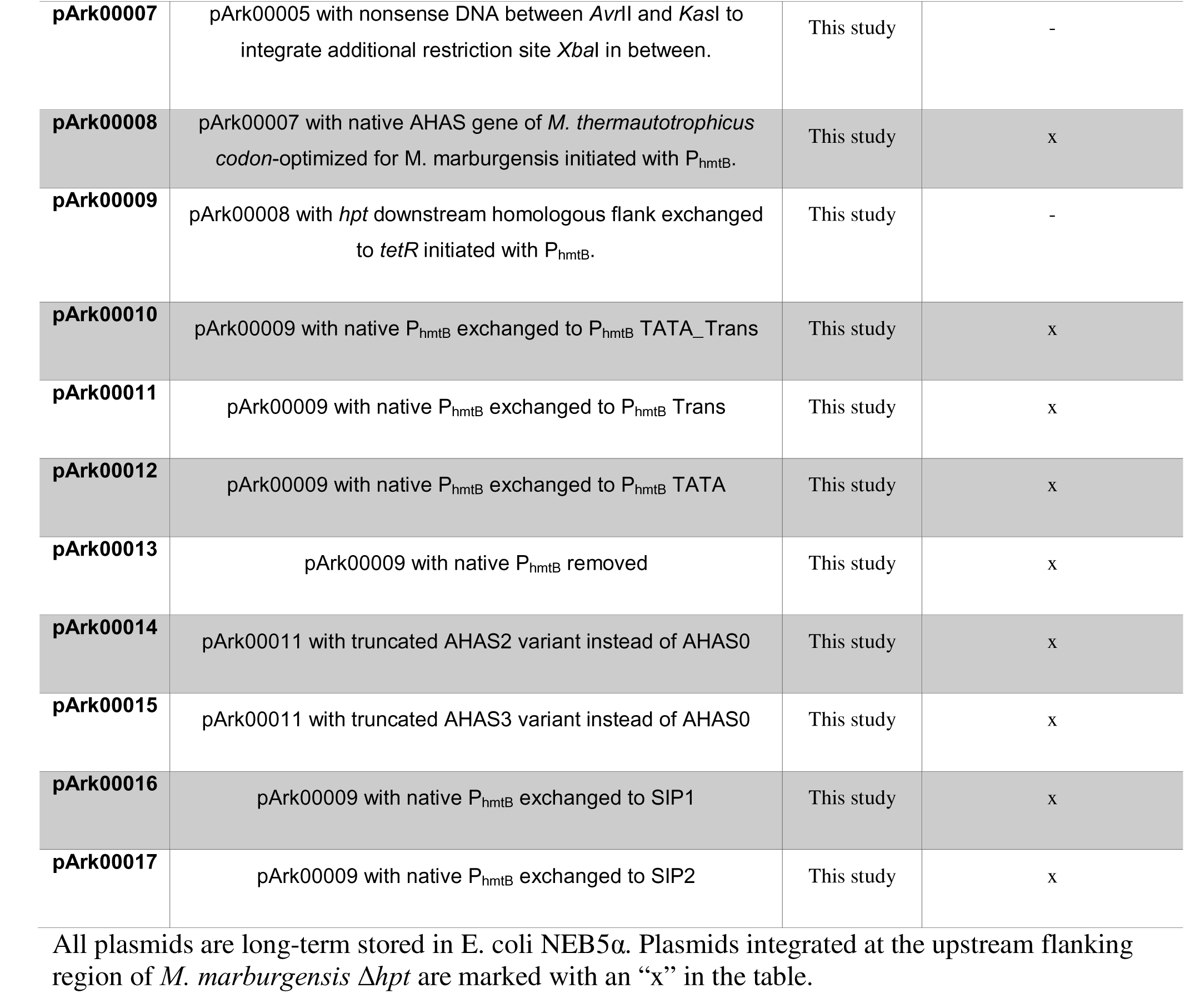
Plasmid list.

**Table 2.**
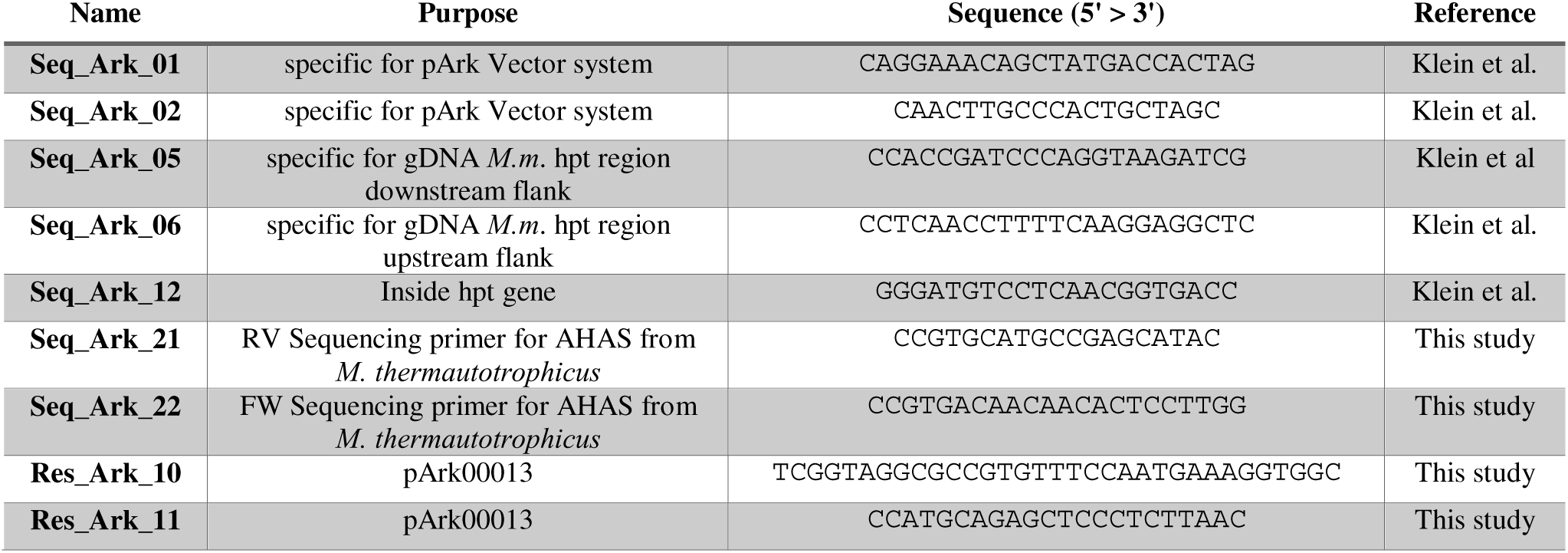
Primer list.

### *E. coli* strains and cultivation

*Escherichia coli* NEB5α from New England Biolabs (New England Biolabs, Ipswich (MA), USA) was used for every cloning step and storage purposes. For the conjugation of *M. marburgensis* with *E. coli*, we utilized the plasmid-mobilizing *E. coli* S17-1 (DSM 9097), purchased from *Deutsche Sammlung für Mikroorganismen und Zellkulturen* (DSMZ, Braunschweig, Germany) (Simon et al. 1983). Both strains were cultivated in liquid Luria Bertani broth (LB broth) (Carl Roth GmbH + Co. KG, Karlsruhe, Germany) at 37 °C with 180 rpm in a shaker incubator (Labwit, Burwood East (VIC), Australia). For selection, 1000-time stocks of chloramphenicol (30 mg mL^-1^), ampicillin (100 mg mL^-1^), or trimethoprim (10 mg mL^-1^) were prepared in ethanol, water and dimethylsulfoxide respectively and kept at -20 °C until use. For solidified media plates, 1.5% (w/v) of Kobe Agar (Carl Roth) was added to the LB broth. After seeding, plates were incubated at 37 °C in a static incubator (Binder GmbH, Tuttlingen, Germany).

### *M. marburgensis* strains, cultivation and transformation

*M. marburgensis* Δ*hpt* was used as the base strain in this study and referenced as wild type throughout the manuscript (Klein et al. 2025). We cultivated *M. marburgensis* under the same conditions as described in (Fink et al. 2021). For conjugation and outgrowth experiments, we used the sea salt media (MS media) formulation. It contained (per liter) sodium chloride, 0.45 g; sodium hydrogen carbonate, 6.00 g; dipotassium hydrogen phosphate, 0.17 g; potassium dihydrogen phosphate, 0.23; ammonium chloride, 0.19 g; magnesium chloride hexahydrate, 0.08 g; calcium chloride dihydrate, 0.06 g; NiCl_2_, 1 ml (0.02 w/v); iron(II)chloride pentahydrate, 1 mL (0.2 w/v); resazurin indicator solution, 4 mL (0.025 w/v); and trace element solution, 1 ml. For amino acid quantifications and enzyme activity assays, *M. marburgensis* was grown in *M. marburgensis* media composition (Abdel Azim et al. 2017; Klein et al. 2025). All gases were purchased with a purity of 5.0 from Air Liquide, Paris, France. Both media compositions were dissolved, the pH adjusted, sparged with N_2_/CO_2_ (20% Vol.-% CO_2_ in N_2_) to remove molecular oxygen (O_2_) and reduced with 0.5 mol L^-1^ Na_2_S·9H_2_O and dispensed under anoxic conditions in an anaerobic chamber (Coy Laboratory Products, Grass Lake (MI), USA) á 20 mL in serum bottles. Serum bottles were then individually cycled through vacuumed and H_2_/CO_2_ (20 Vol.-% CO_2_ in H_2_) atmosphere. The final atmosphere was left with the latter mix at a pressure of 2 bar overpressure. All media contained 4 mL L^-1^ of a 0.025% (w/v) resazurin sodium salt solution as a redox indicator. For selective growth of genetically modified strains, 250 µg mL^-1^ of neomycin was supplemented.

We added 1.5 % Bacto™ agar (Becton, Dickinson and Company, Franklin Lakes (NJ), USA) to solidified media plates and 2.5 g L^-1^ of peptone and 1.25 g L^-1^ of yeast extract to conjugation plates for spot mating.

Transformation of *M. marburgensis* was carried out via conjugal DNA transfer from *E. coli* S17-1 to *M. marburgensis* with spot mating as previously described in Klein et al. 2025. *M. marburgensis* and plasmid carrying *E. coli* S17 were cultivated to late exponential growth phase. Then, ∼2·10^8^ cells of *M. marburgensis* and ∼2·10^9^ *E. coli* cells were centrifuged anaerobically and both pellets were combined in sterile anaerobic MS medium inside the anaerobic chamber. The cell suspension was then carefully poured as spots on conjugation plates prepared as described above. The spots were left to be absorbed until dry and then we incubated under 1 bar overpressure of H_2_/CO_2_ (20 Vol.-% CO_2_ in H_2_) in a stain-less steel jar for 18 h at 37 °C. Afterwards, spots were washed and dissolved in 1 mL of sterile MS media. The transformants were then recovered in 5 mL of MS media for 4 h at 62 °C. 1 mL of recovered *M. marburgensis* was subsequently transferred to 20 mL of neomycin-containing MS medium for liquid selection of positive transformants. After confirmation of the presence of the right mutant via PCR, monoclonal transformants were picked from selective purity plating. Then, the mutants were confirmed and ready for further tests. The PCR followed a colony PCR-like protocol following the Phire plant mastermix manufactureŕs manual (Thermo Scientific, Waltham (MA), USA) (https://phaidra.univie.ac.at/detail/o:2120011).

### Phenotypic characterization of *M. marburgensis* strains

The standard phenotypic characterization was performed in *M. marburgensis* media. A serum bottle was inoculated with an overnight culture incubated at 62 °C to reach a start OD_600_ of 0.05 (n=3). The cultures were incubated at 62 °C over the course of 24 h and samples for amino acid and optical density measurement were taken every 2 h or 3 h and 24 h after the inoculation. For comparison between two incubation temperatures this protocol was adapted. First, the overnight culture was cultivated at 55 °C. In the morning, the culture was cooled down to room temperature. A small volume was taken to inoculate a fresh media bottle from the same batch to OD_600_ of 0.05. The grown culture was then regased with H_2_/CO_2_ to 2 bars and incubated at 62 °C and the OD_600_ of 0.05 bottle was incubated at 55 °C. The sampling procedure was performed in the same manner.

### Anoxic enzymatic activity assay of AHAS variants

For the enzyme assay, cultures were grown in *M. marburgensis* media. Then 4.5 mL of an OD_600_=0.3 culture was centrifuged in an anaerobic chamber (Coy Laboratory Products, Grass Lake (MI), USA). The pellet is resuspended in 500 μL sterile ultrapure water and beadbeated with 0.5 mm zirconium beads two times 20 s with 20 s with 20 s of break in between (Fastprep-24, MP Biomedicals, Irvine, CA, USA). To separate zirconium beads and cell debris, we centrifuged the mix 5 min, 13000 g, at room temperature and anaerobically transferred the supernatant to a sterile reaction tube. All the steps involving the enzymatic reactions were done anaerobically in the anaerobic chamber (Xing and Whitman 1987).

The assay buffer was composed of 20 mmol L^-1^ sodium pyruvate, 0.5 mmol L^-1^ Thiamine diphosphate, 10 μmol L^-1^ Flavin adenine dinucleotide, and 10 mmol ^-1^ MgCl_2_ in 50 mmol ^-1^ Tris buffer (50 mmmol L^-1^ Tris-HCl, 150 mmol ^-1^ NaCl, pH=7.0). It was anaerobized with N_2_/CO_2_ for 30 min on ice. The buffer was prepared fresh, kept on ice and used the same or following day. We mixed 20 µL of assay buffer with 2 µL of *M. marburgensis* crude extract in 0.5 mL PCR tubes (ThermoFisher Scientific, Waltham, MA, USA). The tubes were anaerobically incubated for 1 h at 60 °C. Thereafter, we added 2 µL of 4 mol L^-1^ sulfuric acid, mixed well, and incubated at 60 °C aerobically to convert acetolactate to the more stable acetoin by spontaneous decarboxylation. To develop the characteristic red color for the colorimetric assay, we mixed creatine (0.5% (w/v)) and naphthol (5% naphthol (w/v) in 2.5 mol L^-1^ NaOH) in a 1:1 ratio and added 36 µL of the mixture to the sample. After further vortexing and incubation at 60 °C for 15 min, we centrifuged the samples to remove any crude extract precipitate for 5 min, 13000 g, at room temperature and measured the supernatant absorbance at 525 nm in 384-well plate (Greiner Bio, Kremsmünster, AT) with a Tecan Spark plate reader (Tecan Group Ltd, Maennedorf, Switzerland). To measure the allosteric inhibition of AHAS by valine, leucine and isoleucine, we added 0 to 25 mmol L^-1^ of valine, leucine, or isoleucine, respectively, to the assay mix. This was done by making a twofold concentrated assay buffer and diluting it with solutions of twice the concentration of BCAA. For each enzyme, results are shown as ratios of the initial activity as the starting activity varies greatly.

### Amino acid and acetoin quantification

In total 20 proteinogenic amino acids and acetoin as decarboxylated stable product of acetolactate as valine biosynthesis pathway intermediate were quantified directly from the supernatant of *M. marburgensis* cultures. For amino acid quantification, after the removal of cells, the supernatant was diluted in acetonitrile (1:3) and warmed up for 5 minutes to 40 °C to facilitate protein precipitation. After centrifugation (14 000 g; 5 min), the supernatant was mixed with the IS (4:1; IS was 10 µmol L^-1^ of Metabolomics Amino Acid Mix purchased from Cambridge Isotope Laboratories, Inc.). For the chromatographic separation the LC system (Agilent 1260 Infinity II Prime LC) was coupled to a MS detector (Agilent 6475 Triple Quadrupole) and separation was achieved with a Poroshell 120 HILIC-Z analytical column (2.7 µm, 2.1x100 mm) together with gradient elution from 90/10 to 65/35 25 mmol L^-1^ aqu. NH_4_COO/MeCN (pH=3). Analytes were ionized using ESI in positive mode and analyzed with dynamic MRM. A 9-point external calibration curve with concentrations from 76 nmol L^-1^ to 125 µmol L^-1^ was used for quantification. For analysis of acetoin and other intermediates, samples were derivatized with 3-nitrophenylhydrazine. In short, samples were mixed with IS containing citric acid and pyruvic acid. For derivatization 50% ethanol solutions containing 3-nitrophenylhydrazine, 1-ethyl-3-carbodiimide and pyridine were added and samples were shaken at room temperature for 60 minutes. Analysis was carried out on the same LC/MS system using Poroshell 120 EC-C18 analytical column (2.7 µm, 2.1x100 mm) and a gradient elution from 0.1% formic acid to 90% acetonitrile. ESI in positive mode was used for ionization coupled with dynamic MRM. For quantification a 6-point external calibration curve ranging from 32 nmol L^-1^ to 100 µmol L^-1^ was used.

### Bioinformatics

We performed a structural analysis of *M. thermautotrophicus* AHAS in comparison to AHAS from *E. coli* and *C. glutamicum*. We used the representative AlphaFold structures (Varadi et al. 2022) of the AHAS enzymes from Uniprot and aligned them in PyMOL (The PyMOL Molecular Graphics System, Version 3.0 Schrödinger, LLC.) using the *align* function, highlighted respective valine binding pockets and colored the enzyme structures to facilitate the differentiation. For the correspondence between positions in the sequence of the gene, a multisequence alignment was generated in Snapgene using the MUSCLE algorithm. To check for the correct alignment, we made sure that the mutations that were found to be in the same location in the AHAS of *E. coli* and *C. glutamicum* were correctly aligned, which was the case. We used this alignment to deduce which amino acids to change in *M. thermautotrophicus* AHAS.

### Data interpretation

Different lengths of lag phase might affect the growth stage of the microbe. To compensate for that, triplicates and other multiplets were normalized at each time point and for each strain (**Equation 1**). This accounts for small variations in OD, as the growth phase at similar times should be comparable. Of course, in case outliers were detected, they were left out from the analysis.

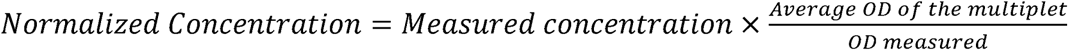

Equation 1. Normalization of amino acid concentration.

The normalized data of the amino acid concentrations built the base for further calculations, such as the volumetric productivity or specific productivity. It requires at least 2 timepoint measurements and is defined as shown in **Equation 2**.

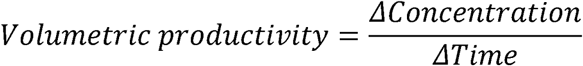

Equation 2. Calculation of volumetric productivity

The specific productivity is then defined as shown in **Equation 3**:

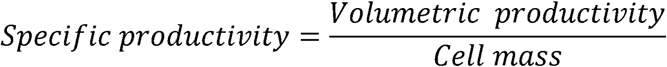

Equation 3. Calculation of specific productivity

We chose to approximate the cell mass as the median point between the starting cell mass and the final cell mass. This underestimates the average cell mass and could slightly overestimate the specific productivity in exponential phase. So, cell mass is defined as shown in **Equation 4** with *a* the factor to convert OD_600_ in grams of cell dry weight equal to 0.371 (Duboc et al. 1995).

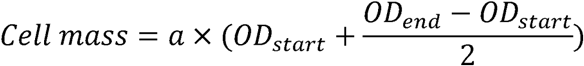

Equation 4. Calculation of average cell mass between time points.

## Results

### TetR DNA-binding affinity regulates temperature-induced valine production in *M. marburgensis*

In the recent years, tools for genetic engineering of *M. marburgensis* have been made available. Those tools were based on promoters for constitutive gene expression. However, to better understand the impact of heterologous gene products on the physiology of *M. marburgensis*, the development of an inducible promoter system was the logical next step. Therefore, we amended the temperature-inducible promoter system based on the tetracycline repressor (TetR) from *E. coli* to *M. marburgensis* which has previously successfully been demonstrated in thermophilic bacteria with similar growth temperature (Mol 2022). This is possible due to the growth rate of *M. marburgensis* being minor impacted by a decrease of 10 °C in incubation temperature around its optimal growth temperature (Ding et al. 2010).

One of the commonly used constitutive promoters for *M. marburgensis* genetic engineering is P_hmtB_ which natively initiates gene expression of a histone-binding protein-encoding gene of *M. thermautotrophicus* ΔH (Klein et al. 2025; Darcy et al. 1999). We chose P_hmtB_ due to its well-understood architecture as a first target for integration of the TetR operator region (TetO2). We designed three putatively inducible promoters in a similar fashion as it has been done for P*_mcrB_* in *M. acetivorans* (Guss et al. 2008). They differ in the localization of TetO2, an 18-bp DNA sequence originally found in *E. coli* (Bertram and Hillen 2008). The resulting P_hmtB_ variants are named according to the position of TetO2, either between the TATA box and transcription start site (TATA), between transcription start site and ribosome binding site (Trans), or both (TATA_Trans) (**Figure 1**).

**Figure 1.**
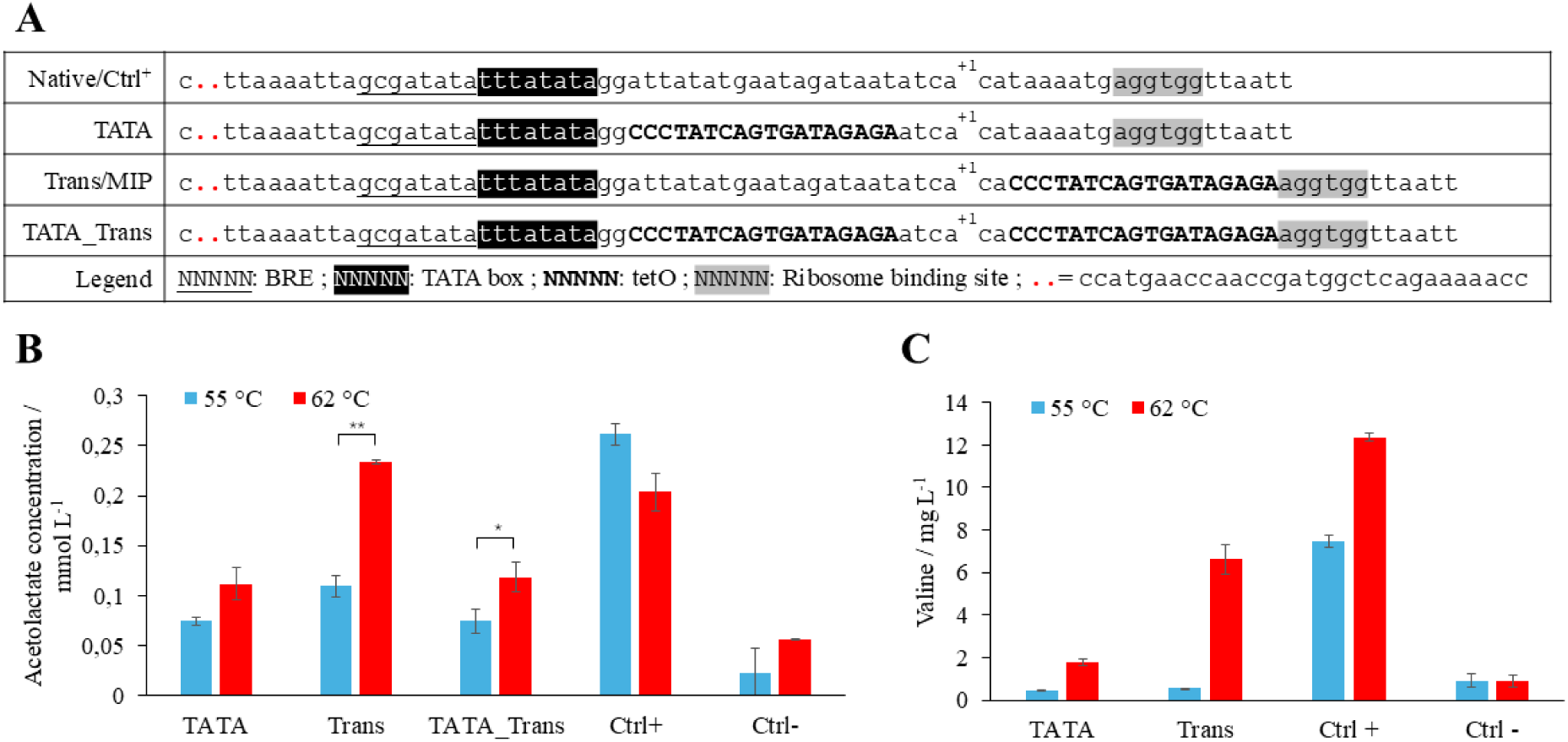
Structure and influence of different putative inducible promoter variants based on P_hmtB_. **A)** Structures of the different promoter variants. The known elements are shown with different formatting defined in the “Legend” row; the B recognition element (BRE), TetR DNA-binding site (TetO2), and the transcription start site (+1). **B)** Acetolactate concentration measured in the AHAS activity assay for each of the *M. marburgensis* mutants containing native AHAS and promoter variants described above at the different temperatures. 55 °C is the “OFF” state and 62 °C is the “ON” state. Ctrl^-^ is the strain with no promoter upstream of AHAS. The error bars indicate the standard deviation (n=3). Asterisks show statistical significance derived from pairwise comparisons in Student’s T-tests (*: p ≤ 0.05; **: p ≤ 0.01). **C)** Valine concentration measured in the supernatant after overnight incubation at the indicated temperatures for each strain presented above. Ctrl^-^ is the strain with no promoter upstream of AHAS. The error bars indicate the standard deviation (n=2).

As a reporter gene for valine production, we positioned the P_hmtB_ variants upstream of the acetohydroxyacid synthase (AHAS)-encoding gene from *M. thermautotrophicus* ΔH codon-optimized for *M. marburgensis.* We utilized the gene of *M. thermautotrophicus* to avoid off-target effects in homologous recombination at the native AHAS-encoding gene of *M. marburgensis*. AHAS marks the entry enzyme into the valine biosynthesis pathway from pyruvate (Huang et al. 2017). This has been shown to result in overproduction of valine in previous studies. Additionally, we added TetR-encoding gene under constitutive expression of P_hmtB_ in the opposite reading direction downstream of the AHAS encoding gene. The resulting construct contained a 1 kbp upstream homologous flanking region of the deleted hypoxanthine phosphoribosyl transferase (*hpt*) of *M. marburgensis* for targeted genome integration and a thermostable neomycin resistance for positive selection in *M. marburgensis* (**Supplementary figure S1**).

First, we generated the respective *M. marburgensis* mutant strains with the three different TetO2-containing P_hmtB_ promoter (TATA, Trans, and TATA_Trans), the native P_hmtB_ (Ctrl^+^), and a mutant with the AHAS gene but without an upstream promoter (Ctrl^-^) (**Table 1**). Afterwards, we performed a temperature screening to gather initial insight into the functionality of the promoter variants, the valine production, and the inducibility thereof. TetR was expected to release from the DNA at around 60 °C (Wagenhöfer et al. 1988; Mol 2022). Therefore, we chose a temperature range between 55 °C and 66 °C. We found increased valine production starting at a temperature of 57 °C with a trend confirmed at temperatures above 60 °C (**Supplementary figure S2**). Since our goal is to apply the temperature-inducible promoter system to large-scale bioreactor processes, the temperature change needs to remain as small as possible. Therefore, we chose the temperature differences of 7 °C between 55 °C (OFF state) and 62 °C (ON state) as our standard operation temperature in bioreactors even if stronger effects were visible at 66 °C (**Supplementary figure S3**).

After the screening approach, we aimed for a more precise assessment of the activity during the ON and OFF states of each promoter variant to decide which one was most suitable for further studies. In the first step, we performed a AHAS enzyme activity assay on crude extract of the *M. marburgensis* mutant strain with the respective promoter variants (TATA, Trans, TATA_Trans, Ctrl^+^,Ctrl^-^) (**Figure 1B**). While we observed a trend towards higher activity in the ON state compared to the OFF state in TATA and TATA_Trans (p=0.05 to p=0.1), we found a significant increase in activity in Trans (p ≤ 0.01). It reaches similar activity levels as AHAS in the Ctrl^+^ in ON state and only around 43% of activity compared to its OFF state. However, we still observed a residual activity of AHAS in crude extract from the Trans mutant in the Off state compared to the basal activity of Ctrl^-^. To measure the effect of the different promoters on actual valine production, the valine concentration was measured from an overnight culture of the respective *M. marburgensis* mutant strains in the OFF state and in the ON state (**Figure 1C**). Between the OFF and ON states, we measured a 1.6-fold increase in valine concentration in the Ctrl^+^. The valine concentration of TATA and Trans, however, showed a 4-fold and 12-fold increase in the ON state, respectively. Furthermore, the valine concentration in the OFF state of the TATA and Trans mutant is comparably low as in the basal activity of Ctrl^-^ (**Figure 1C**). In conclusion, we showed that the temperature-inducible promoter system based on TetR is functional in *M. marburgensis,* with corresponding results of the AHAS enzyme activity assay and the valine concentration in the supernatant. It appears that the Trans promoter variant is superior in terms of tightness and expression levels compared to the other variants tested. Therefore, we chose the Trans variant for subsequent studies and renamed it to Medium Inducible Promoter (MIP).

### Integration of TetO2 in P_glnA(*M.v*.)_ increases regulatory effect and valine secretion in *M. marburgensis* compared to MIP

With the temperature induction proven in MIP, we wanted to further increase the valine production rates in the ON state while decreasing the basal activity in the OFF state. In a promoter study of four native promoters which are commonly used in other methanogen genetic systems, we identified in reporter gene assays higher activity with P_glnA(*M.v*.)_ from *Methanococcus vannielli* compared to P_synth(BRE)_ and P_hmtB_ (**Supplementary figure S4**) (Fink et al. 2021). In contrast to the constitutive promoter P_hmtB_, P *_glnA(M.v)_* natively contains an operator region due to its nitrogen-regulated characteristic (Bao et al. 2022). To not interfere with the general promoter architecture of P *_glnA(M.v)_*, we exchanged this operator region with TetO2. Additionally, we generated a second promoter variant with TetO2 upstream of the ribosome binding site as it was done in MIP. We named those promoters Strong Inducible Promoter (SIP) 2 and 1, respectively (**Figure 2A**).

**Figure 2.**
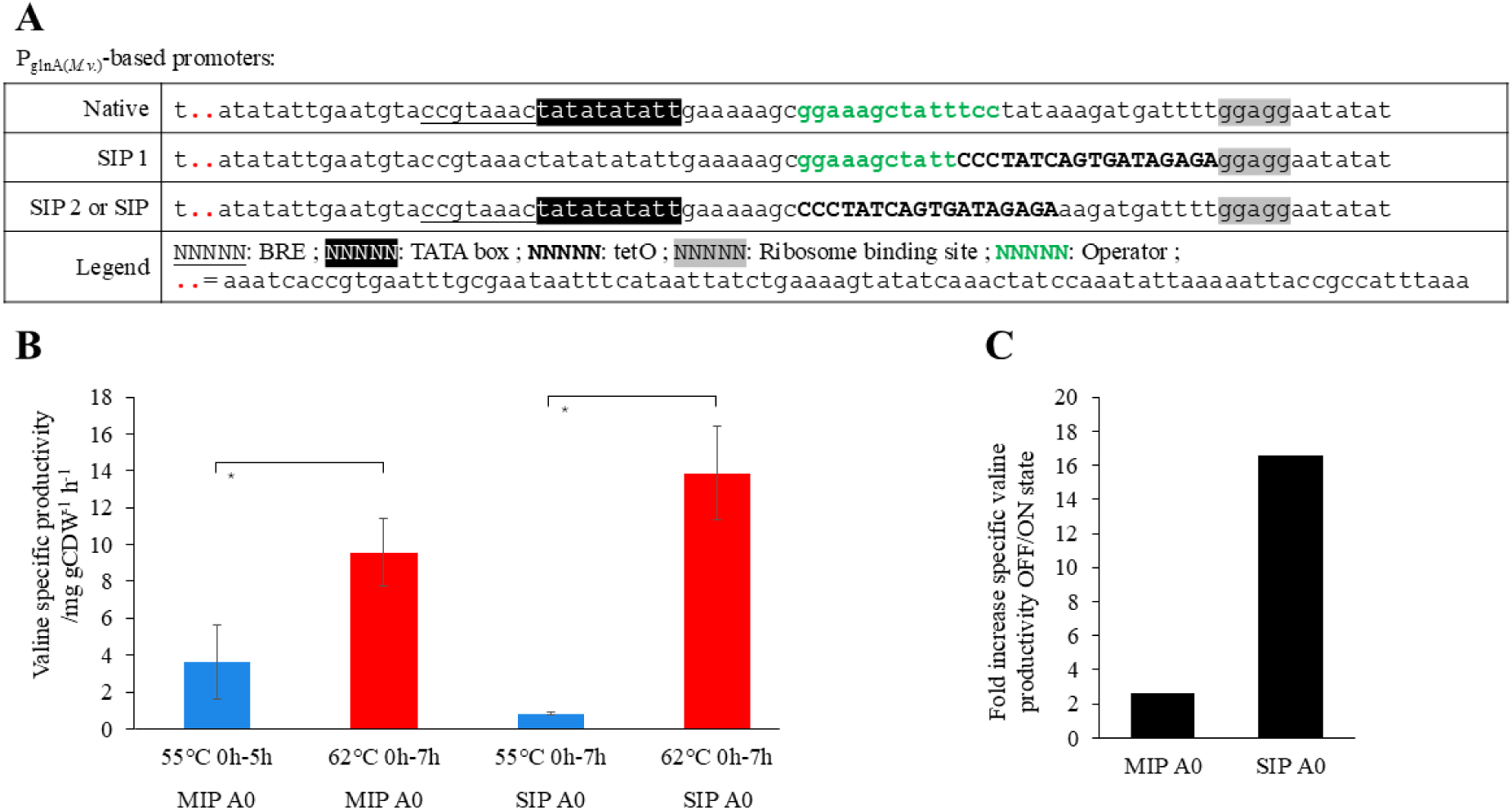
Structure and influence of different putative inducible promoters based on P_glnA(*M.v*.)_. **A**) Structures of the different promoters. The different known elements are shown with different formatting which are explained in the “Legend” row. BRE stands for B recognition element, TetO2 for tetracycline operator. **B**) Valine specific production rate during exponential phase at the indicated temperatures for each strain presented above. The error bars indicate the standard deviation (n=3). Paired Student’s T-Test for comparison of OFF and ON state with p<0.05 marked with asterisk. **C**) Fold increase of the specific valine productivity between 55 °C (OFF state) and 62 °C (ON state). Ratio of averages. (n=3)

In a first screening approach (n=1) with SIP1 and SIP2 containing *M. marburgensis* mutants upstream of AHAS, we found SIP1 to have a 2.3-fold increase in valine concentration in the supernatant between the OFF and ON states and a 21.5-fold increased valine concentration with SIP2 (**Supplementary figure S5**). Therefore, we continued our study with the SIP2 containing *M. marburgensis* mutant strain. The phenotypic characterization (n=3) confirmed the results from the initial screening. Here, we demonstrated a 12.9-fold increase in specific valine productivity between the OFF state and ON state for SIP2, while under the same conditions MIP only showed a 1.5-fold increase (**Figure 2C**). Additionally, the specific valine productivity in the ON state during the exponential growth phase was found to be significantly higher in SIP2 with 14.17 mg gCDW^-1^ h^-1^ compared to MIP with 9.01 mg gCDW^-1^ h^-1^ (**Figure 2B**). Thus, we conclude that SIP2 initiates tighter temperature-induced expression paired with stronger gene expression that results in higher specific valine production rates than MIP.

### Release of allosteric valine inhibition in AHAS enhances specific valine production rates in *M. marburgensis*

Another path to follow to engineer microbial cell factories for valine overproduction is the release of allosteric feedback inhibition from the acetolactate synthase (AHAS) enzyme involved in valine biosynthesis. In the case of AHAS, it was shown to be allosteric inhibited by the downstream BCAAs and especially valine (Elišáková et al. 2005). The AHAS enzymes are always assembled from two dimers, one from the large (catalytic) subunit and one from the small (regulatory) subunit. However, the regulatory subunit also influences the functionality of the catalytic subunit. Therefore, removal or deletion of the small subunit is not an option for the release of allosteric inhibition. It is actually a strategy to decrease activity of AHAS to improve the production of other compounds (Blombach et al. 2009). A multisequence alignment of the amino acid sequences of the small regulatory subunit of AHAS from *E. coli*, *C. glutamicum*, and *M. thermautotrophicus* showed a strong conservation of the regulatory subunit of AHAS amongst bacteria and archaea (**Figure 3A**). Especially in the region of the confirmed valine binding pocket for allosteric inhibition of the enzyme in *E. coli* and *C. glutamicum*, AHAS from *M. thermautotrophicus* has a high level of conservation (**Figure 3A**).

**Figure 3.**
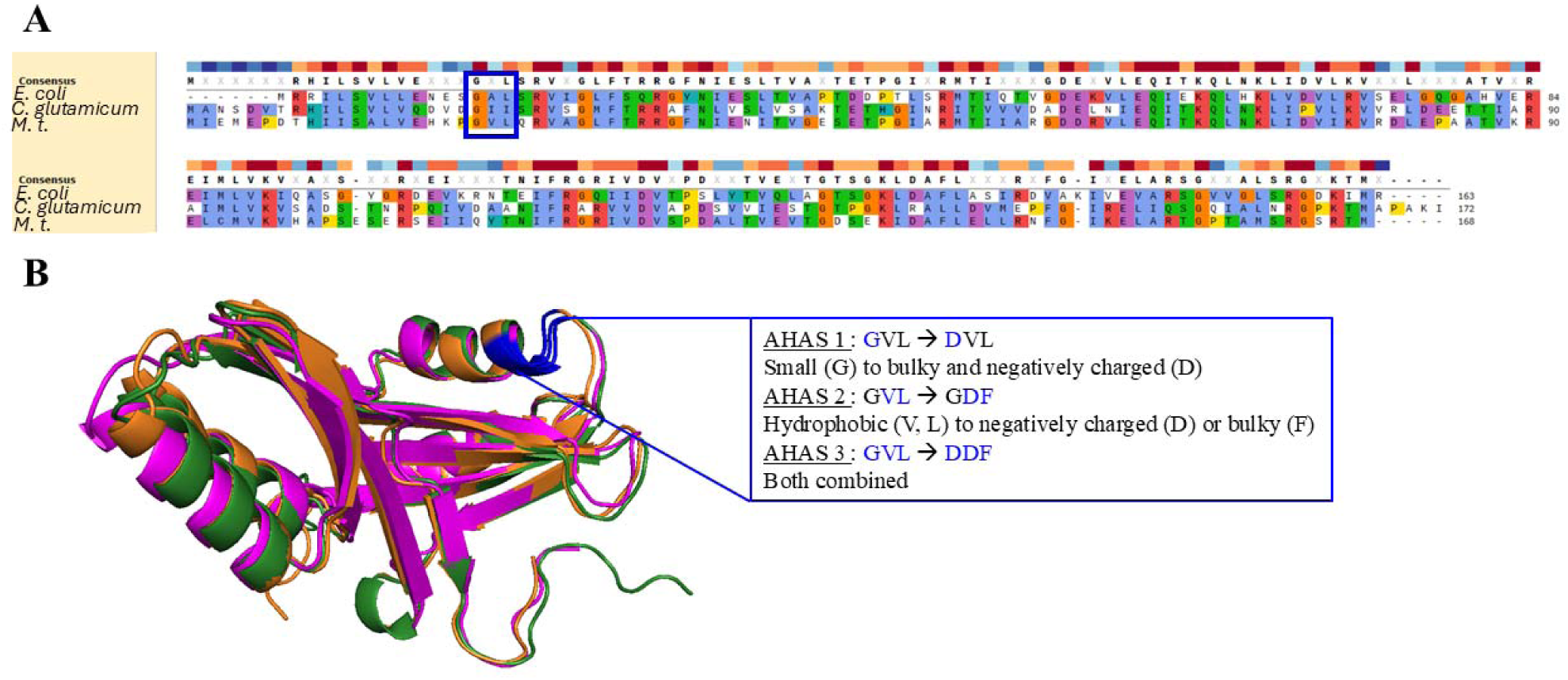
Design of inhibition released AHAS variant enzymes. **A**) Multisequence alignment of native AHAS amino acid sequences of *E. coli*, *C. glutamicum*, and *M. thermautotrophicus* (*M.t*.). **B**) Alignment of the structure of the same enzymes. Green, magenta and orange proteins represent AHAS from *C. glutamicum*, *E. coli* and *M. thermautotrophicus* respectively. Residues in blue were the ones mutated. Box shows mutations done on each mutant of the AHAS from *M. thermautotrophicus*. Letters are standard one-letter code for amino acids by IUPAC.

In two conclusive review and descriptive publications about the regulatory AHAS subunit of *E. coli* (Kaplun et al. 2006; Elišáková et al. 2005), the impact of spontaneous mutagenized versions of AHAS and rationally designed mutant versions was tested on allosteric resistance. In Kaplun *et al*. a collection of rationally designed mutant variants and experimental tests thereof was given including the differences in valine inhibition. The G14 (red residue on **Figure 3**) and following amino acids in *E. coli* seem to have the greatest impact on allosteric inhibition through valine. Alpha-fold structures of the *E. coli* small subunit compared to *M. thermautotrophicus* small subunit appear relatively similar in terms of the whole structure of the enzyme but also in the respective area of G14 upstream of helix 1 (**Figure 3B**). Based on the literature research, we integrated the crucial mutations in AHAS to remove allosteric inhibition in *E. coli* and *C. glutamicum* and designed three different variants of the AHAS from *M. thermautotrophicus*, such as I) AHAS 1, G20D (A1), II) AHAS 2 (A2), V21D, L22F, and III) AHAS 3 (A3): G20D, V21D, L22F. In the same pattern, the wild-type enzyme is also referred to as AHAS 0 (A0). We did not manage to successfully transform *M. marburgensis* with the A1 variant in several approaches. Therefore, we characterized the A2 and A3 variants only.

The *M. marburgensis* mutants of A2 and A3 were then generated with the MIP (P_hmtB__Trans) promoter. Mutations in regulatory regions of enzymes can lead to loss or decrease of function. So, the first thing we evaluated was the activity of the AHAS variants without inhibitory substances, such as valine or leucine. A2 variant maintained 69% of the wild type A0 variant activity, whereas the A3 variant showed only 25% of the A0 activity (**Figure 4A**). Furthermore, the enzyme assay supported and strengthened our findings on the inducibility of the MIP promoter, with an increase in activity of around 10% compared to wild type in the OFF state (55 °C) compared to basal activity.

**Figure 4.**
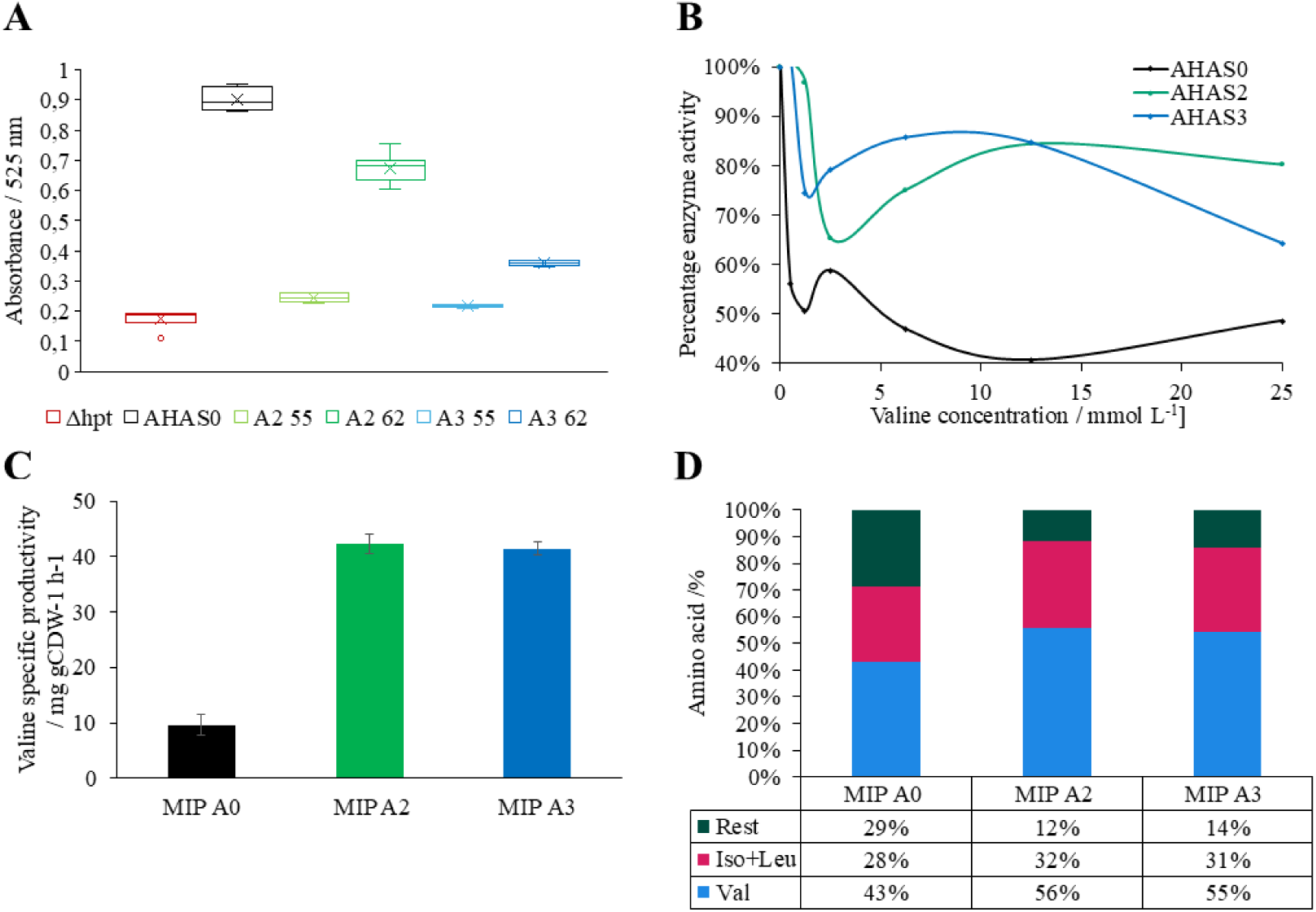
Effects on specific valine production for different allosteric inhibition released AHAS variants (AHAS0, AHAS2, AHAS3). **A)** Relative activity of each strain cultivated at different temperatures shown as the OD_600_ obtained after the assay. *M. marburgensis* wild-type represents the base strain from which all the others tested derive. AHAS0 contains the AHAS from *M. thermautotrophicus*. A2 (AHAS2) and A3 (AHAS3) have the putative inhibition released enzymes. 55 and 62 indicate the growth temperatures in degree Celsius. (n=6) **B)** Activity of each AHAS variant with different valine concentrations from 0 to 25 mmol L^-1^ as allosteric inhibitor (n=2) **C)** Valine specific productivity during exponential phase at the indicated temperatures for each strain. The error bars indicate the standard deviation (n=3). **D)** Amino acid ratio after overnight growth for *M. marburgensis* mutant strains with AHAS0 (A0), AHAS2 (A2), and AHAS3 (A3) integrated and expressed with medium strength inducible promoter (MIP) (n=3).

To test the activity at high valine concentrations, the enzymatic AHAS assay was adapted to test increasing valine concentrations from 0-25 mmol L^-1^. The trend shows a quick collapse of the activity of the wild-type A0 down to around 50 to 40% of the original activity at 2.5 mmol L^-1^ of valine. A3 and especially A2 hold their activity much better with up to 80% of the initial activity even at higher valine concentrations (**Figure 4B, Supplementary figure S6**). To elucidate the influence of the inhibition release regarding the specific valine productivity, we performed a phenotypic characterization (n=3). It shows an increase in specific valine productivity from 10.7 mg gCDW^-1^ h^-1^ in A0 to 57.6 mg gCDW^-1^ h^-1^ and 17.8 mg gCDW^-1^ h^-1^ for A2 and A3 respectively. This represents almost a 6-fold increase in specific valine production rates (**Figure 4C**). Additionally, it is reflected in the ratio of valine compared to total amino acids obtained. While the valine content compared to all 20 proteinogenic amino acids in the supernatant was 43% on average in A0, we found a significant increase to 56, 55% in A2 and A3 respectively and an increase from 71% to 88% (A2) and 86% (A3) for BCAAs (**Figure 4D**). All enzymes show at least some residual activity, so the specific productivity of valine was measured. (**Figure 4C**). A2 and A3 show a 4.4- and 4.3-fold increase in specific productivity respectively compared to allosteric inhibited A0.

With the release of inhibition of one enzyme in an enzyme cascade, the product of the reaction might accumulate if the downstream enzymes are not effective enough. In our case, the chemical compound likely to accumulate would be acetolactic acid. It is the product of the AHAS enzymatic reaction. Our acetolactic acid measurement in the supernatant (**Supplementary figure S7**) showed a 15-18-fold increase.

## Discussion

In this study, we successfully amended a temperature-controlled inducible promoter system for *M. marburgensis* based on the TetR system and applied it for inducible overproduction of the proteinogenic amino acid valine. After proof of concept with the commonly used non-regulated constitutive P_hmtB_ promoter for *M. marburgensis*, we further refined the induction capacity and promoter strength by the integration of the TetR binding site (TetO2) in the P_glnA(*M.v*.)_ promoter from *Methanococcus vannielli.* In a second approach, we achieved valine overproduction by removal of the putative allosteric valine binding site in the acetolactate synthase (AHAS) enzyme from *M. thermautotrophicus* based on alpha-fold model structural comparisons to AHAS structures from *E. coli* and *C. glutamicum*. The exchanges of up to three amino acids in the respective valine binding region resulted in reduced sensitivity towards allosteric valine inhibition and significantly higher specific production rates of valine of up to 4-fold compared to the wild-type AHAS overexpression mutant of *M. marburgensis*.

In the amendment of the TetR inducible promoter system for *M. marburgensis*, we first used the constitutive promoter P_hmtB_ as a basis, which is functional in *M. thermautotrophicus* and *M. marburgensis* but also shows a large host range since it is also utilized in *Thermococcus kodakarensis* (Fink et al. 2021; Klein et al. 2025; Santangelo et al. 2008). The well-characterized P_hmtB_ promoter architecture from BRE element, via TATA box, to transcription start and ribosome binding site facilitated the choice of where to integrate the TetR binding site without interference with essential parts of the promoter (Darcy et al. 1999). The placement of the TetR binding site was performed analogously to the integration sites in the P_mcrB_ promoter for usage in *Methanosarcina acetivorans,* which is up to now the only methanogen genus where the TetR system was shown to be functional (Guss et al. 2008). Integration of the TetR binding site in P_hmtB_ at a similar position in combination with a betagalactosidase enzyme assay yielded anhydrotetracycline (aTc)-induced gene expression in *M. thermautotrophicus.* There, it has been shown that the aTc concentration has a significant influence on the induction of gene-expression (Baur et al. 2025). The P_mcrB_ promoter follows the same architecture as the P_hmtB_ promoter which facilitated the identification of the integration sites (Zhu et al. 2024). In *E. coli*, there are two TetR binding sites described, TetO1 and TetO2. In coherence with literature, we chose TetO2 as it was shown to result in tighter repression and stronger inducibility of the gene of interest compared to TetO1 (Bertram and Hillen 2008). In terms of the position of TetO2, we found the position between transcription start site and ribosome binding site (Trans) as the most suitable for P_hmtB_. The ON state was not affected and inducibility could be proven with the OFF state. The position between TATA box and transcription start site resulted in reduced activity during the ON state (**Figure 1B**). This is exactly the opposite as described for P_mcr_ in *M. acetivorans* (Guss et al. 2008). The fluctuation of results in regard to the position of TetO2 seems to be dependent on the promoter architecture used as a basis for gene expression in the respective host (Bertram and Hillen 2008).

The TetR repressor is released from TetO1 and TetO2 using tetracycline or anhydrotetracycline due to its reduced antimicrobial impact on bacterial and archaeal cells (Bertram et al. 2022, Baur et al. 2025). However, it has been demonstrated that the TetR repressor detaches from its binding sites at ∼60 °C (Wagenhöfer et al. 1988). This fact makes it suitable for thermo-induction for thermophilic microbes with a growth range between 50 and 70 °C. In studies with *Parageobacillus thermoglucosidasius*, the TetO1 binding site has been used for thermo-induction where below 60 °C was defined as the lower cutoff point (Mol 2022). We found inducibility already from 57 °C with TetO2 as TetR binding site and P_hmtB_ as promoter basis forming a gradient up to 65 °C (**Supplementary figure S3**). To use the thermo-induction in industrial-scale processes, the delta between the OFF and ON states needs to be as small as possible due to heating costs of the bioreactor systems and the time loss for production during the heating process. For a 100 m³ reactor that would represent 814 kWh which is already a vast amount of energy to put in a running reactor quickly for rapid induction. (Isobaric specific heat of water at 55 °C: 0.00116 kWh (kg K)^-1^ according to which is 814 kWh for a 100 000 kg of water). Therefore, we chose 55 °C as the OFF state and 62 °C as the ON state with the risk of not operating at full induction (**Supplementary figure S3**). The use of TetO1 instead of TetO2 as TetR binding site could offer benefits in the future since the delta between OFF and ON state appears smaller and a certain leakiness of the induction could be used to generate product already during the growth stage without high burden on the metabolism (Mao et al. 2024).

To circumvent the limits of induction at 62 °C and for a tighter gene expression, we further developed the TetR thermo-induction system by the use of P_glnA(*M.v*.)_ from *M. vannielli* as promoter basis. The promoter architecture resembles the one from P_hmtB_ and is fully characterized (Bao et al. 2022). The only difference is the operator region for ammonia and alanine induction discovered in *M. maripaludis* (Lie et al. 2005). This operator region fitted TetO2 well with only 2 bp difference in length. Therefore, we exchanged the native operator for TetO2. Additionally, the promoter strength has been shown to be relatively strong compared to other constitutive promoters in *M. maripaludis* (Bao et al. 2022). We also measured higher activity of betagalactosidase in enzymatic assays with P_glnA(*M.v*.)_ compared to P_hmtB_ or P_synth_ (Fink et al. 2021) (**Supplementary figure S4**). With the integration of thermo-inducible P_glnA(*M.v*.)_, we enhanced to the tightness to wild-type-like levels and increased the specific valine productivity by 5.5-fold compared to the P_hmtB_ based inducible promoter (Trans) (**Figure 2B**).

In our second approach to enhance valine production rates and concentration, we aimed for removal of allosteric valine inhibition in the *M. thermautotrophicus* AHAS enzyme. We found the AHAS enzyme structure in *M. thermautotrophicus* to be highly homologous compared to *E. coli* or *C. glutamicum* (**Figure 3A+B**) (Cordes et al. 1992; Park and Lee 2010). This homology also included the allosteric valine binding pocket that has been proven in *E. coli* and *C. glutamicum*. To prevent valine from binding this pocket and therefore release allosteric inhibition, up to three amino acids have been exchanged in that region of the enzyme. This resulted in complete release of valine inhibition in AHAS enzymes from *E. coli*, *C. glutamicum*, and *Streptomyces cinnamonensis* (Elišáková et al. 2005; Kopecký et al. 1999). We implemented the same amino acid exchanges in heterologous produced AHAS enzymes in *M. marburgensis* and indeed found reduced sensitivity to valine as well (**Figure 4B**). It demonstrates that mutations across domains of life with strong homology have the same effect. However, as has been observed in *M. maripaludis* already, also AHAS from *M. thermautotrophicus* is highly O_2_ sensitive, which is not the case in well-characterized AHAS enzymes from *E. coli* or *C. glutamicum* (Xing and Whitman 1987). With the modified AHAS variants from *M. thermautotrophicus* expressed in *M. marburgensis*, we found a 4-fold increased specific valine productivity compared to mutants overexpressing the wild-type AHAS enzyme. The 40 mg gCDW^-1^ h^-1^ specific valine productivity is in a similar range as the reported valine productivities in *C. glutamicum* and *E. coli* with similar genetic modifications of 5 to 110 mg gCDW^-1^ h^-1^ (Elišáková et al. 2005; Gao et al. 2021). Recently, also the recombinant production of acetoin with *M. thermautotrophicus* has been reported. Since acetoin production and valine production share the AHAS enzyme in the pathway, the biosynthesis is highly comparable. In closed-batch cultivation 0.17 mg OD_600_ ^-1^ L^-1^ h^-1^ specific productivity has been demonstrated (Baur et al. 2025) This fact confirms that *M. marburgensis* presents a suitable platform for commodity chemical production. In the next step, we will test the valine overproduction strains of *M. marburgensis* in a fed-batch bioreactor set up and try to scale the specific productivity to reach higher volumetric productivity, since higher cell densities are possible in reactor systems (up to 8 / g L^-1^) compared to 0.08 / g L^-1^ in closed batch experiments (Rittmann et al. 2018; Hofmann et al. 2025). Hence, bioreactor cell densities of *M. marburgensis* in fed-batch processes can reportedly match cell densities for *C. glutamicum* and *E. coli* in common reactor systems for valine production (Du et al. 2025; Buchholz et al. 2013).

Another great advantage when using methanogens for commodity chemical production, such as BCAAs, is the low byproduct formation. With methane gassing out of the media, the supernatant of closed batch experiments for valine production contained up to 88% BCAA. To eliminate the share of BCAAs other than valine, such as isoleucine and leucine, it was already shown that the deletion of the citramalate synthase (*cimA*) leads to isoleucine auxotrophy (Klein et al. 2025). The same holds true for leucine via isopropylmalate synthase (*leuA*) deletion that has been shown in *M. maripaludis* (Hendrickson et al. 2008). Afterwards, a bradytrophic strain for isoleucine and leucine could be generated as has been shown for *E. coli* (Park et al. 2007). Additional room for further development leaves the optimization of the carbon flux from pyruvate to valine as we found intermediates such as acetolactic acid to accumulate in the supernatant of valine overproduction strains of *M. marburgensis*. One possibility to enhance the carbon flux represents fine-tuning and overexpression of downstream genes which harbor great potential for substantially increased volumetric and specific production rates of valine (Gao et al. 2021). Furthermore, we experienced a substantial decrease of enzyme activity in the truncated AHAS variants in *M. marburgensis*. This appears typical when modifying AHAS enzymes and is consistent with findings of Elišáková et al. 2005 and offers great potential for further development of the AHAS enzyme variants with the help of protein engineering (Tan et al. 2024).

In conclusion, we showed that with the integration of state-of-the-art tools, such as protein engineering and a thermos-inducible promoter, *M. marburgensis* shows the capability for overproduction of valine with relatable specific production rates common hosts for sugar-based production of valine. This marks the next advance towards commodity chemical production with methanogenic archaea. As the scale-up of bioreactor cultivation with *M. marburgensis* has been shown before Haslinger, Reischl et al. 2025, submitted for publication). A combination of a further developed microbial cell factory of *M. marburgensis* with adapted bioprocess conditions for commodity chemical production will provide the basis for *M. marburgensis* as a suitable platform for large-scale product formation.

## Author Contributions

C.F., R.F., and S.K.-M. R. R. initiated the project. C.F. and R.F. planned the operational tasks. S.M., F.U., and M.K. performed the laboratory experiments. T.S. did analytical measurements. S.M., F.U. and C.F. analysed the data. R.F., C.F. and J.S. supervised the project. F.U., C.F., and S.K.-M. R. R. wrote the manuscript. S.K.-M. R. R. acquired funding. All authors edited the manuscript and approved its publication.

## Funding

The COMET center: acib: Next Generation Bioproduction is funded by BMIMI, BMWET, SFG, Standortagentur Tirol, Government of Lower Austria und Vienna Business Agency in the framework of COMET - Competence Centers for Excellent Technologies. The COMET-Funding Program is managed by the Austrian Research Promotion Agency FFG. Open access funding was provided by the University of Vienna.

## Acknowledgments

The authors thank Angus Hilts for the helpful conversations about bioinformatic analyses. We also thank Stefan Drießler for his input on cloning experiments and laboratory management.

## Data Availability Statement

The raw data set for this study can be found on the PHAIDRA University of Vienna data repository with the following DOI: 10.25365/phaidra.724.

## Notes

### Competing Interest Statement

The authors have declared no competing interest.

