## Supplementary Material for "Valine overproduction with metabolically engineered *Methanothermobacter marburgensis*"

### 1 Supplementary Methods

#### ONPG assay

To assess promoter strength of various promoters in *Methanothermobacter marburgensis*, each of them were ligated to a betagalactosidase reporter encoding gene using overlap extension PCR (Fink et al. 2021). Promoters were synthetised as gBlocks ordered from IDT genomics. Then, the promoter-Bgab fragments were digested and ligated into an integration vector and transformed in *M. marburgensis*. We cultivated each promoter and betagalactosidase (BgaB) containing strain overnight. Afterwards, the pellet of each sample (3 mL of $OD_{600}$=0.3) was resuspended in 85 µl *M. marburgensis* media and beat beated for 3x20 s. After the lysis the samples were centrifuged for 5 min at 21300 g to pellet the debris. The supernatant was used for the assay a technical triplicate of each sample created by pipetting three times 25 µL of each sample into respective wells of a 96-well plate. As a negative control we used sterile *M. marburgensis* media and wild-type *M. marburgensis* and tested in the assay as a technical triplicate of each. 175 µL of ONPG reaction solution was added to each well on top of the crude extract and mixed via vortexing (Fink et al. 2021; Jensen et al. 2017). The plate was sealed with a breathable foil and incubated at 65 °C. The first measurement at 420 nm absorbance was performed after 30 mins in a Tecan Sunrise plate reader. Additionally, measurements were performed after 60, 90, and 120 min. There were no air bubbles present at all measurements. Based on the measurements, the respective Miller units were calculated using the formula Absorbance [420 nm]/(Incubation time [h] · $OD_{600}$ · Volume [L]).

Geometric means and standard deviations were calculated for each sample using the calculated Miller unit values of the technical triplicates. Afterwards bar charts were generated.

### 2 Supplementary Results

#### Synthesized fragments

**Supplementary table S1**: Synthesized fragments

| Type | Name | Insert ordered | DNA delivered |
|---|---|---|---|
| Gene | AHAS0 codon optimized | TCTAGAAAATGTCCTGAAATAAAAAAAATTAGGTGCATTACACTTAAAAA<br>GTTTGTGCAATGCACCTTCGTTAATCAGTATCGGCTCACATTGTCCTGCTT<br>CCCCTGGACATTGCTGTTGGCCCTGTCCTTGCGAGTTCCTTTATTCCAAAG<br>TTCCTCAGAAGCTCCAGGAAGGCGTCTATCTTTTCTGAGTCCCCTGTAACC<br>TCAACTGTCAGGGCATCGGGTGACACATCCACTATCCTCCCCCTGAATATG<br>TTGGTGTACTGTATTATCTCTGACCTTTCTGACTCTGATGGGGCGTGGACC<br>TTGACCATGCAGAGCTCCCTCTTAACTGTTGCTGCGGGTTCAAGGTCCCTG<br>ACCTTTATGACGTCTATGAGCTTGTTCAGCTGCTTTGTTATCTGCTCAAGC<br>ACCCTGTCGTCGCCCCTGGCGATTATTGTCATCCTGGCAATGCCTGGGGTT<br>TCTGATTCCCCGACTGTTATGTTTTCAATGTTGAATCCCCTTCTTGTGAAGA<br>GTCCTGCAACCCTCTGGAGCACTCCTGGTTTGTGCTCCACGAGGGCGCTTA<br>TGATATGGGTATCGGGTTCCATCTCAATCACCATCCGCCTTCCCTGGGGAG<br>TATTTAATTTCCTGGGGGTCCTCCCTTTCAACCCTGTACTCCCCCACTATCT<br>CGGTGAGACCACAGCCGGGGGGGACCATGGGGAGTATCTCATCAGGATCT<br>ATCACTATATCAAGGAGGGCAGGTTCACCTGACCTTATTGCCCTTGAAAG<br>GGCTTCTGAGGTTTCACCGGGTTCCTCTATCCTCTCTGCCTCCACTCCAAA<br>TGATTCTGCCAGCTTCACAAAGTCGGGAACTTCGCCCAGGTGTGTATGGG<br>ACATTCTCTCATCATAGAAGAGCCTCTGCCACTGTGCCACCATTCCAAGGT<br>GCCTGTTGTCCATGATACATATCACCACGGGGATGTCGTATTCCCTTATGG<br>TTGCAAGGTCCTGGCAGACCATGAGGAATCCGCCGTCACCGCACACTGCA<br>ACAACGTCTGAATCAGGCAGTGCCACCTTGGCACCTATGGCGGCTGGAAA<br>ACCGAAGCCCATTGTTCCAAGGCCTCCTGATGATATGAACTTTCTGGGGG<br>CCCTGGATGTGTAGAAGTGGGCCATCCACATCTGGTTCTGTCCCACATCTG<br>TTGTAACGACTGTCTCGTCATCAAGGACCTGGCTTATCTCCTTTATAACCT<br>GCTGGGGCTTCAGGGGCACCTCATCATAGCTCATCCTTGGCATGCAATCG<br>GCCCTGAATTTCTGGACGCTTTCAAGCCACTGGCTGTCCCTCTTTTCATATT<br>TTTTGAGTTTTGCTATGAGTTCCCTGAGGACGTTTCTTGCATCTCCAACGA<br>TGGGGACATCAACCCCAACGTTCTTACCTATCTCTGCGGGGTCGATGTCGA<br>CGTGTATTATCCTGGCGTTGGGGGCGAATTCTGCAACGTTCCCTGTTGTCC<br>TGTCTGAGAATCTGCATCCAACGGCTATGAGGCAGTCGCATTCGTCCACT<br>GTCAGGTTTGCCACCTTCCTGCCGTGCATGCCGAGCATACCCATGGCTGAA<br>GGGTGGTCCTCAGGAAAGGAACCCTTACCAAGGAGTGTTGTTGTCACGGG<br>GGCCTTTATGAGATCTGAGAGTTCCTTTATCTCCCTGGATGCCCCTGATAT<br>TATAACTCCTCCACCTGCAAGTATGACGGGTTTTTCTGACCTCCTTATGAG<br>TTCTGCGGCCCTCTTTATCTGGAGGGGGTGGCCCTTAACATTGGGCCTGTA<br>CCCTGGGAGCTCCAGGTCATCAACCTCCTCCATGATCTCCTGTTCCTGTAT<br>ATCCTTGGGGAGGTCTATAACAACGGGTCCTGGCCTTCCTGTCTTTGCTAT<br>GTGGAAGCTTGCCCTGACAATTGCAGGTATCTCGCTGGCGTCTGATGGCT<br>GGAATGAGTGCTTGGTGATGGGCATGGTTATCCCTATCATGTCCACCTCCT<br>GGAATGCATCATTTCCAATGAGGTGTGTTGGGACCTGACCTGCAATGGCC<br>ACGATGGGGGCTGAGTCCATGTAGGCTGTTGCAATGCCTGTAACAAGGTT<br>TGTTGCCCCGGGACCGGAGGTTGCTATGCAGACCCCCACCCTTCCTGAGG<br>CCCTTGCATATCCGTCTGCTGCGTGTGCTGCGCACTGTTCATGTCTAACGA<br>GGATGTGTTTAAGTTCTGAATCATAGAGCATATCATAGAGTGGCAGGAGC<br>TGTCCACCGGGGTATCCGAAAACGGTGTCTGCTCCCTGATCCAGAAGTGA | TCTAGAAAATGTCCTGAAATAAAAAAAATTAGGTGCATTACACTTAAAAA<br>GTTTGTGCAATGCACCTTCGTTAATCAGTATCGGCTCACATTGTCCTGCTT<br>CCCCTGGACATTGCTGTTGGCCCTGTCCTTGCGAGTTCCTTTATTCCAAAG<br>TTCCTCAGAAGCTCCAGGAAGGCGTCTATCTTTTCTGAGTCCCCTGTAACC<br>TCAACTGTCAGGGCATCGGGTGACACATCCACTATCCTCCCCCTGAATATG<br>TTGGTGTACTGTATTATCTCTGACCTTTCTGACTCTGATGGGGCGTGGACC<br>TTGACCATGCAGAGCTCCCTCTTAACTGTTGCTGCGGGTTCAAGGTCCCTG<br>ACCTTTATGACGTCTATGAGCTTGTTCAGCTGCTTTGTTATCTGCTCAAGC<br>ACCCTGTCGTCGCCCCTGGCGATTATTGTCATCCTGGCAATGCCTGGGGTT<br>TCTGATTCCCCGACTGTTATGTTTTCAATGTTGAATCCCCTTCTTGTGAAGA<br>GTCCTGCAACCCTCTGGAGCACTCCTGGTTTGTGCTCCACGAGGGCGCTTA<br>TGATATGGGTATCGGGTTCCATCTCAATCACCATCCGCCTTCCCTGGGGAG<br>TATTTAATTTCCTGGGGGTCCTCCCTTTCAACCCTGTACTCCCCCACTATCT<br>CGGTGAGACCACAGCCGGGGGGGACCATGGGGAGTATCTCATCAGGATCT<br>ATCACTATATCAAGGAGGGCAGGTTCACCTGACCTTATTGCCCTTGAAAG<br>GGCTTCTGAGGTTTCACCGGGTTCCTCTATCCTCTCTGCCTCCACTCCAAA<br>TGATTCTGCCAGCTTCACAAAGTCGGGAACTTCGCCCAGGTGTGTATGGG<br>ACATTCTCTCATCATAGAAGAGCCTCTGCCACTGTGCCACCATTCCAAGGT<br>GCCTGTTGTCCATGATACATATCACCACGGGGATGTCGTATTCCCTTATGG<br>TTGCAAGGTCCTGGCAGACCATGAGGAATCCGCCGTCACCGCACACTGCA<br>ACAACGTCTGAATCAGGCAGTGCCACCTTGGCACCTATGGCGGCTGGAAA<br>ACCGAAGCCCATTGTTCCAAGGCCTCCTGATGATATGAACTTTCTGGGGG<br>CCCTGGATGTGTAGAAGTGGGCCATCCACATCTGGTTCTGTCCCACATCTG<br>TTGTAACGACTGTCTCGTCATCAAGGACCTGGCTTATCTCCTTTATAACCT<br>GCTGGGGCTTCAGGGGCACCTCATCATAGCTCATCCTTGGCATGCAATCG<br>GCCCTGAATTTCTGGACGCTTTCAAGCCACTGGCTGTCCCTCTTTTCATATT<br>TTTTGAGTTTTGCTATGAGTTCCCTGAGGACGTTTCTTGCATCTCCAACGA<br>TGGGGACATCAACCCCAACGTTCTTACCTATCTCTGCGGGGTCGATGTCGA<br>CGTGTATTATCCTGGCGTTGGGGGCGAATTCTGCAACGTTCCCTGTTGTCC<br>TGTCTGAGAATCTGCATCCAACGGCTATGAGGCAGTCGCATTCGTCCACT<br>GTCAGGTTTGCCACCTTCCTGCCGTGCATGCCGAGCATACCCATGGCTGAA<br>GGGTGGTCCTCAGGAAAGGAACCCTTACCAAGGAGTGTTGTTGTCACGGG<br>GGCCTTTATGAGATCTGAGAGTTCCTTTATCTCCCTGGATGCCCCTGATAT<br>TATAACTCCTCCACCTGCAAGTATGACGGGTTTTTCTGACCTCCTTATGAG<br>TTCTGCGGCCCTCTTTATCTGGAGGGGGTGGCCCTTAACATTGGGCCTGTA<br>CCCTGGGAGCTCCAGGTCATCAACCTCCTCCATGATCTCCTGTTCCTGTAT<br>ATCCTTGGGGAGGTCTATAACAACGGGTCCTGGCCTTCCTGTCTTTGCTAT<br>GTGGAAGCTTGCCCTGACAATTGCAGGTATCTCGCTGGCGTCTGATGGCT<br>GGAATGAGTGCTTGGTGATGGGCATGGTTATCCCTATCATGTCCACCTCCT<br>GGAATGCATCATTTCCAATGAGGTGTGTTGGGACCTGACCTGCAATGGCC<br>ACGATGGGGGCTGAGTCCATGTAGGCTGTTGCAATGCCTGTAACAAGGTT<br>TGTTGCCCCGGGACCGGAGGTTGCTATGCAGACCCCCACCCTTCCTGAGG<br>CCCTTGCATATCCGTCTGCTGCGTGTGCTGCGCACTGTTCATGTCTAACGA<br>GGATGTGTTTAAGTTCTGAATCATAGAGCATATCATAGAGTGGCAGGAGC<br>TGTCCACCGGGGTATCCGAAAACGGTGTCTGCTCCCTGATCCAGAAGTGA |

| | | | |
|---|---|---|---|
| | | TCTGATTATTGCCTGGCCACCTTTCATTGGAAACACAATTAACCACCTCAT<br>TTTATGTGATATTATCTATTCATATAATCCTATATAAATATATCGCTAATTT<br>TAAGGTTTTTCTGAGCCATCGGTTGGTTCATGGGGGCGCC | TCTGATTATTGCCTGGCCACCTTTCATTGGAAACACAATTAACCACCTCAT<br>TTTATGTGATATTATCTATTCATATAATCCTATATAAATATATCGCTAATTT<br>TAAGGTTTTTCTGAGCCATCGGTTGGTTCATGGGGGCGCCAGGCTAGGTG<br>GAGGCTCAGTGATGATAAGTCTGCGATGGTGGATGCATGTGTCATGGTCA<br>TAGCTGTTTCCTGTGTGAAATTGTTATCCGCTCAGAGGGCACAATCCTATT<br>CCGCGCTATCCGACAATCTCCAAGACATTAGGTGGAGTTCAGTTCGGCGT<br>ATGGCATATGTCGCTGGAAAGAACATGTGAGCAAAAGGCCAGCAAAAGG<br>CCAGGAACCGTAAAAAGGCCGCGTTGCTGGCGTTTTTCCATAGGCTCCGC<br>CCCCCTGACGAGCATCACAAAAATCGACGCTCAAGTCAGAGGTGGCGAA<br>ACCCGACAGGACTATAAAGATACCAGGCGTTTCCCCCTGGAAGCTCCCTC<br>GTGCGCTCTCCTGTTCCGACCCTGCCGCTTACCGGATACCTGTCCGCCTTT<br>CTCCCTTCGGGAAGCGTGGCGCTTTCTCATAGCTCACGCTGTAGGTATCTC<br>AGTTCGGTGTAGGTCGTTCGCTCCAAGCTGGGCTGTGTGCACGAACCCCC<br>CGTTCAGCCCGACCGCTGCGCCTTATCCGGTAACTATCGTCTTGAGTCCAA<br>CCCGGTAAGACACGACTTATCGCCACTGGCAGCAGCCACTGGTAACAGGA<br>TTAGCAGAGCGAGGTATGTAGGCGGTGCTACAGAGTTCTTGAAGTGGTGG<br>CCTAACTACGGCTACACTAGAAGAACAGTATTTGGTATCTGCGCTCTGCTG<br>AAGCCAGTTACCTTCGGAAAAAGAGTTGGTAGCTCTTGATCCGGCAAACA<br>AACCACCGCTGGTAGCGGTGGTTTTTTTGTTTGCAAGCAGCAGATTACGCG<br>CAGAAAAAAAGGATCTCAAGAAGATCCTTTGATCTTTTCTACGGGGTCTG<br>ACGCTCTATTCAACAAAGCCGCCGTCCCGTCAAGTCAGCGTAAATGGGTA<br>GGGGGCTTCAAATCGTCCTCGTGATACCAATTCGGAGCCTGCTTTTTTGTA<br>CAAACTTGTTGATAATGGCAATTCAAGGATCTTCACCTAGATCCTTTTAAA<br>TTAAAAATGAAGTTTTAAATCAATCTAAAGTATATATGAGTAAACTTGGT<br>CTGACAGTTACCAATGCTTAATCAGTGAGGCACCTATCTCAGCGATCTGTC<br>TATTTCGTTCATCCATAGTTGCCTGACTCCCCGTCGTGTAGATAACTACGA<br>TACGGGAGGGCTTACCATCTGGCCCCAGTGCTGCAATGATACCGCGAGAG<br>CCACGCTCACCGGCTCCAGATTTATCAGCAATAAACCAGCCAGCCGGAAG<br>GGCCGAGCGCAGAAGTGGTCCTGCAACTTTATCCGCCTCCATCCAGTCTAT<br>TAATTGTTGCCGGGAAGCTAGAGTAAGTAGTTCGCCAGTTAATAGTTTGC<br>GCAACGTTGTTGCCATTGCTACAGGCATCGTGGTGTCACGCTCGTCGTTTG<br>GTATGGCTTCATTCAGCTCCGGTTCCCAACGATCAAGGCGAGTTACATGAT<br>CCCCCATGTTGTGCAAAAAAGCGGTTAGCTCCTTCGGTCCTCCGATCGTTG<br>TCAGAAGTAAGTTGGCCGCAGTGTTATCACTCATGGTTATGGCAGCACTG<br>CATAATTCTCTTACTGTCATGCCATCCGTAAGATGCTTTTCTGTGACTGGT<br>GAGTACTCAACCAAGTCATTCTGAGAATAGTGTATGCGGCGACCGAGTTG<br>CTCTTGCCCGGCGTCAATACGGGATAATACCGCGCCACATAGCAGAACTT<br>TAAAAGTGCTCATCATTGGAAAACGTTCTTCGGGGCGAAAACTCTCAAGG<br>ATCTTACCGCTGTTGAGATCCAGTTCGATGTAACCCACTCGTGCACCCAAC<br>TGATCTTCAGCATCTTTTACTTTCACCAGCGTTTCTGGGTGAGCAAAAACA<br>GGAAGGCAAAATGCCGCAAAAAAGGGAATAAGGGCGACACGGAAATGTT<br>GAATACTCATACTCTTCCTTTTTCAATATTATTGAAGCATTTATCAGGGTT<br>ATTGTCTCATGAGCGGATACATATTTGAATGTATTTAGAAAAATAAACAA<br>ATAGGGGTTCCGCGCACATTTCCCCGAAAAGTGCCAGATACCTGAAACAA<br>AACCCATCGTACGGCCAAGGAAGTCTCCAATAACTGTGATCCACCACAAG<br>CGCCAGGGTTTTCCCAGTCACGACGTTGTAAAACGACGGCCAGTCATGCA<br>TAATCCGCACGCATCTGGAATAAGGAAGTGCCATTCCGCCTGACCT |
| Gene | AHAS2 | TCTAGAAAATGTCCTGAAATAAAAAAAATTAGGTGCATTACACTTAAAAA<br>GTTTGTGCAATGCACCTTCGTTAATCAGTATCGGCTCACATTGTCCTGCTT<br>CCCCTGGACATTGCTGTTGGCCCTGTCCTTGCGAGTTCCTTTATTCCAAAG<br>TTCCTCAGAAGCTCCAGGAAGGCGTCTATCTTTTCTGAGTCCCCTGTAACC<br>TCAACTGTCAGGGCATCGGGTGACACATCCACTATCCTCCCCCTGAATATG<br>TTGGTGTACTGTATTATCTCTGACCTTTCTGACTCTGATGGGGCGTGGACC<br>TTGACCATGCAGAGCTCCCTCTTAACTGTTGCTGCGGGTTCAAGGTCCCTG<br>ACCTTTATGACGTCTATGAGCTTGTTCAGCTGCTTTGTTATCTGCTCAAGC<br>ACCCTGTCGTCGCCCCTGGCGATTATTGTCATCCTGGCAATGCCTGGGGTT<br>TCTGATTCCCCGACTGTTATGTTTTCAATGTTGAATCCCCTTCTTGTGAAGA<br>GTCCTGCAACCCTCTGGAAATCACCTGGTTTGTGCTCCACGAGGGCGCTTA<br>TGATATGGGTATCGGGTTCCATCTCAATCACCATCCGCCTTCCCTGGGGAG<br>TATTTAATTTCCTGGGGGTCCTCCCTTTCAACCCTGTACTCCCCCACTATCT<br>CGGTGAGACCACAGCCGGGGGGGACCATGGGGAGTATCTCATCAGGATCT<br>ATCACTATATCAAGGAGGGCAGGTTCACCTGACCTTATTGCCCTTGAAAG<br>GGCTTCTGAGGTTTCACCGGGTTCCTCTATCCTCTCTGCCTCCACTCCAAA<br>TGATTCTGCCAGCTTCACAAAGTCGGGAACTTCGCCCAGGTGTGTATGGG<br>ACATTCTCTCATCATAGAAGAGCCTCTGCCACTGTGCCACCATTCCAAGGT<br>GCCTGTTGTCCATGATACATATCACCACGGGGATGTCGTATTCCCTTATGG<br>TTGCAAGGTCCTGGCAGACCATGAGGAATCCGCCGTCACCGCACACTGCA | TCTAGAAAATGTCCTGAAATAAAAAAAATTAGGTGCATTACACTTAAAAA<br>GTTTGTGCAATGCACCTTCGTTAATCAGTATCGGCTCACATTGTCCTGCTT<br>CCCCTGGACATTGCTGTTGGCCCTGTCCTTGCGAGTTCCTTTATTCCAAAG<br>TTCCTCAGAAGCTCCAGGAAGGCGTCTATCTTTTCTGAGTCCCCTGTAACC<br>TCAACTGTCAGGGCATCGGGTGACACATCCACTATCCTCCCCCTGAATATG<br>TTGGTGTACTGTATTATCTCTGACCTTTCTGACTCTGATGGGGCGTGGACC<br>TTGACCATGCAGAGCTCCCTCTTAACTGTTGCTGCGGGTTCAAGGTCCCTG<br>ACCTTTATGACGTCTATGAGCTTGTTCAGCTGCTTTGTTATCTGCTCAAGC<br>ACCCTGTCGTCGCCCCTGGCGATTATTGTCATCCTGGCAATGCCTGGGGTT<br>TCTGATTCCCCGACTGTTATGTTTTCAATGTTGAATCCCCTTCTTGTGAAGA<br>GTCCTGCAACCCTCTGGAAATCACCTGGTTTGTGCTCCACGAGGGCGCTTA<br>TGATATGGGTATCGGGTTCCATCTCAATCACCATCCGCCTTCCCTGGGGAG<br>TATTTAATTTCCTGGGGGTCCTCCCTTTCAACCCTGTACTCCCCCACTATCT<br>CGGTGAGACCACAGCCGGGGGGGACCATGGGGAGTATCTCATCAGGATCT<br>ATCACTATATCAAGGAGGGCAGGTTCACCTGACCTTATTGCCCTTGAAAG<br>GGCTTCTGAGGTTTCACCGGGTTCCTCTATCCTCTCTGCCTCCACTCCAAA<br>TGATTCTGCCAGCTTCACAAAGTCGGGAACTTCGCCCAGGTGTGTATGGG<br>ACATTCTCTCATCATAGAAGAGCCTCTGCCACTGTGCCACCATTCCAAGGT<br>GCCTGTTGTCCATGATACATATCACCACGGGGATGTCGTATTCCCTTATGG<br>TTGCAAGGTCCTGGCAGACCATGAGGAATCCGCCGTCACCGCACACTGCA |

| | | | |
|---|---|---|---|
| | | ACAACGTCTGAATCAGGCAGTGCCACCTTGGCACCTATGGCGGCTGGAAA<br>ACCGAAGCCCATTGTTCCAAGGCCTCCTGATGATATGAACTTTCTGGGGG<br>CCCTGGATGTGTAGAAGTGGGCCATCCACATCTGGTTCTGTCCCACATCTG<br>TTGTAACGACTGTCTCGTCATCAAGGACCTGGCTTATCTCCTTTATAACCT<br>GCTGGGGCTTCAGGGGCACCTCATCATAGCTCATCCTTGGCATGCAATCG<br>GCCCTGAATTTCTGGACGCTTTCAAGCCACTGGCTGTCCCTCTTTTCATATT<br>TTTTGAGTTTTGCTATGAGTTCCCTGAGGACGTTTCTTGCATCTCCAACGA<br>TGGGGACATCAACCCCAACGTTCTTACCTATCTCTGCGGGGTCGATGTCGA<br>CGTGTATTATCCTGGCGTTGGGGGCGAATTCTGCAACGTTCCCTGTTGTCC<br>TGTCTGAGAATCTGCATCCAACGGCTATGAGGCAGTCGCATTCGTCCACT<br>GTCAGGTTTGCCACCTTCCTGCCGTGCATGCCGAGCATACCCATGGCTGAA<br>GGGTGGTCCTCAGGAAAGGAACCCTTACCAAGGAGTGTTGTTGTCACGGG<br>GGCCTTTATGAGATCTGAGAGTTCCTTTATCTCCCTGGATGCCCCTGATAT<br>TATAACTCCTCCACCTGCAAGTATGACGGGTTTTTCTGACCTCCTTATGAG<br>TTCTGCGGCCCTCTTTATCTGGAGGGGGTGGCCCTTAACATTGGGCCTGTA<br>CCCTGGGAGCTCCAGGTCATCAACCTCCTCCATGATCTCCTGTTCCTGTAT<br>ATCCTTGGGGAGGTCTATAACAACGGGTCCTGGCCTTCCTGTCTTTGCTAT<br>GTGGAAGCTTGCCCTGACAATTGCAGGTATCTCGCTGGCGTCTGATGGCT<br>GGAATGAGTGCTTGGTGATGGGCATGGTTATCCCTATCATGTCCACCTCCT<br>GGAATGCATCATTTCCAATGAGGTGTGTTGGGACCTGACCTGCAATGGCC<br>ACGATGGGGGCTGAGTCCATGTAGGCTGTTGCAATGCCTGTAACAAGGTT<br>TGTTGCCCCGGGACCGGAGGTTGCTATGCAGACCCCCACCCTTCCTGAGG<br>CCCTTGCATATCCGTCTGCTGCGTGTGCTGCGCACTGTTCATGTCTAACGA<br>GGATGTGTTTAAGTTCTGAATCATAGAGCATATCATAGAGTGGCAGGAGC<br>TGTCCACCGGGGTATCCGAAAACGGTGTCTGCTCCCTGATCCAGAAGTGA<br>TCTGATTATTGCCTGGCCACCTTTCATTGGAAACACAATTAACCACCTCAT<br>TTTATGTGATATTATCTATTCATATAATCCTATATAAATATATCGCTAATTT<br>TAAGGTTTTTCTGAGCCATCGGTTGGTTCATGGGGGCGCC | ACAACGTCTGAATCAGGCAGTGCCACCTTGGCACCTATGGCGGCTGGAAA<br>ACCGAAGCCCATTGTTCCAAGGCCTCCTGATGATATGAACTTTCTGGGGG<br>CCCTGGATGTGTAGAAGTGGGCCATCCACATCTGGTTCTGTCCCACATCTG<br>TTGTAACGACTGTCTCGTCATCAAGGACCTGGCTTATCTCCTTTATAACCT<br>GCTGGGGCTTCAGGGGCACCTCATCATAGCTCATCCTTGGCATGCAATCG<br>GCCCTGAATTTCTGGACGCTTTCAAGCCACTGGCTGTCCCTCTTTTCATATT<br>TTTTGAGTTTTGCTATGAGTTCCCTGAGGACGTTTCTTGCATCTCCAACGA<br>TGGGGACATCAACCCCAACGTTCTTACCTATCTCTGCGGGGTCGATGTCGA<br>CGTGTATTATCCTGGCGTTGGGGGCGAATTCTGCAACGTTCCCTGTTGTCC<br>TGTCTGAGAATCTGCATCCAACGGCTATGAGGCAGTCGCATTCGTCCACT<br>GTCAGGTTTGCCACCTTCCTGCCGTGCATGCCGAGCATACCCATGGCTGAA<br>GGGTGGTCCTCAGGAAAGGAACCCTTACCAAGGAGTGTTGTTGTCACGGG<br>GGCCTTTATGAGATCTGAGAGTTCCTTTATCTCCCTGGATGCCCCTGATAT<br>TATAACTCCTCCACCTGCAAGTATGACGGGTTTTTCTGACCTCCTTATGAG<br>TTCTGCGGCCCTCTTTATCTGGAGGGGGTGGCCCTTAACATTGGGCCTGTA<br>CCCTGGGAGCTCCAGGTCATCAACCTCCTCCATGATCTCCTGTTCCTGTAT<br>ATCCTTGGGGAGGTCTATAACAACGGGTCCTGGCCTTCCTGTCTTTGCTAT<br>GTGGAAGCTTGCCCTGACAATTGCAGGTATCTCGCTGGCGTCTGATGGCT<br>GGAATGAGTGCTTGGTGATGGGCATGGTTATCCCTATCATGTCCACCTCCT<br>GGAATGCATCATTTCCAATGAGGTGTGTTGGGACCTGACCTGCAATGGCC<br>ACGATGGGGGCTGAGTCCATGTAGGCTGTTGCAATGCCTGTAACAAGGTT<br>TGTTGCCCCGGGACCGGAGGTTGCTATGCAGACCCCCACCCTTCCTGAGG<br>CCCTTGCATATCCGTCTGCTGCGTGTGCTGCGCACTGTTCATGTCTAACGA<br>GGATGTGTTTAAGTTCTGAATCATAGAGCATATCATAGAGTGGCAGGAGC<br>TGTCCACCGGGGTATCCGAAAACGGTGTCTGCTCCCTGATCCAGAAGTGA<br>TCTGATTATTGCCTGGCCACCTTTCATTGGAAACACAATTAACCACCTCAT<br>TTTATGTGATATTATCTATTCATATAATCCTATATAAATATATCGCTAATTT<br>TAAGGTTTTTCTGAGCCATCGGTTGGTTCATGGGGGCGCCAGGCTAGGTG<br>GAGGCTCAGTGATGATAAGTCTGCGATGGTGGATGCATGTGTCATGGTCA<br>TAGCTGTTTCCTGTGTGAAATTGTTATCCGCTCAGAGGGCACAATCCTATT<br>CCGCGCTATCCGACAATCTCCAAGACATTAGGTGGAGTTCAGTTCGGCGT<br>ATGGCATATGTCGCTGGAAAGAACATGTGAGCAAAAGGCCAGCAAAAGG<br>CCAGGAACCGTAAAAAGGCCGCGTTGCTGGCGTTTTTCCATAGGCTCCGC<br>CCCCCTGACGAGCATCACAAAAATCGACGCTCAAGTCAGAGGTGGCGAA<br>ACCCGACAGGACTATAAAGATACCAGGCGTTTCCCCCTGGAAGCTCCCTC<br>GTGCGCTCTCCTGTTCCGACCCTGCCGCTTACCGGATACCTGTCCGCCTTT<br>CTCCCTTCGGGAAGCGTGGCGCTTTCTCATAGCTCACGCTGTAGGTATCTC<br>AGTTCGGTGTAGGTCGTTCGCTCCAAGCTGGGCTGTGTGCACGAACCCCC<br>CGTTCAGCCCGACCGCTGCGCCTTATCCGGTAACTATCGTCTTGAGTCCAA<br>CCCGGTAAGACACGACTTATCGCCACTGGCAGCAGCCACTGGTAACAGGA<br>TTAGCAGAGCGAGGTATGTAGGCGGTGCTACAGAGTTCTTGAAGTGGTGG<br>CCTAACTACGGCTACACTAGAAGAACAGTATTTGGTATCTGCGCTCTGCTG<br>AAGCCAGTTACCTTCGGAAAAAGAGTTGGTAGCTCTTGATCCGGCAAACA<br>AACCACCGCTGGTAGCGGTGGTTTTTTTGTTTGCAAGCAGCAGATTACGCG<br>CAGAAAAAAAGGATCTCAAGAAGATCCTTTGATCTTTTCTACGGGGTCTG<br>ACGCTCTATTCAACAAAGCCGCCGTCCCGTCAAGTCAGCGTAAATGGGTA<br>GGGGGCTTCAAATCGTCCTCGTGATACCAATTCGGAGCCTGCTTTTTTGTA<br>CAAACTTGTTGATAATGGCAATTCAAGGATCTTCACCTAGATCCTTTTAAA<br>TTAAAAATGAAGTTTTAAATCAATCTAAAGTATATATGAGTAAACTTGGT<br>CTGACAGTTACCAATGCTTAATCAGTGAGGCACCTATCTCAGCGATCTGTC<br>TATTTCGTTCATCCATAGTTGCCTGACTCCCCGTCGTGTAGATAACTACGA<br>TACGGGAGGGCTTACCATCTGGCCCCAGTGCTGCAATGATACCGCGAGAG<br>CCACGCTCACCGGCTCCAGATTTATCAGCAATAAACCAGCCAGCCGGAAG<br>GGCCGAGCGCAGAAGTGGTCCTGCAACTTTATCCGCCTCCATCCAGTCTAT<br>TAATTGTTGCCGGGAAGCTAGAGTAAGTAGTTCGCCAGTTAATAGTTTGC<br>GCAACGTTGTTGCCATTGCTACAGGCATCGTGGTGTCACGCTCGTCGTTTG<br>GTATGGCTTCATTCAGCTCCGGTTCCCAACGATCAAGGCGAGTTACATGAT<br>CCCCCATGTTGTGCAAAAAAGCGGTTAGCTCCTTCGGTCCTCCGATCGTTG<br>TCAGAAGTAAGTTGGCCGCAGTGTTATCACTCATGGTTATGGCAGCACTG<br>CATAATTCTCTTACTGTCATGCCATCCGTAAGATGCTTTTCTGTGACTGGT<br>GAGTACTCAACCAAGTCATTCTGAGAATAGTGTATGCGGCGACCGAGTTG<br>CTCTTGCCCGGCGTCAATACGGGATAATACCGCGCCACATAGCAGAACTT<br>TAAAAGTGCTCATCATTGGAAAACGTTCTTCGGGGCGAAAACTCTCAAGG<br>ATCTTACCGCTGTTGAGATCCAGTTCGATGTAACCCACTCGTGCACCCAAC<br>TGATCTTCAGCATCTTTTACTTTCACCAGCGTTTCTGGGTGAGCAAAAACA<br>GGAAGGCAAAATGCCGCAAAAAAGGGAATAAGGGCGACACGGAAATGTT<br>GAATACTCATACTCTTCCTTTTTCAATATTATTGAAGCATTTATCAGGGTT<br>ATTGTCTCATGAGCGGATACATATTTGAATGTATTTAGAAAAATAAACAA<br>ATAGGGGTTCCGCGCACATTTCCCCGAAAAGTGCCAGATACCTGAAACAA |

| | | | |
|---|---|---|---|
| | | | AACCCATCGTACGGCCAAGGAAGTCTCCAATAACTGTGATCCACCACAAG<br>CGCCAGGGTTTTCCCAGTCACGACGTTGTAAAACGACGGCCAGTCATGCA<br>TAATCCGCACGCATCTGGAATAAGGAAGTGCCATTCCGCCTGACCT |
| Gene | AHAS3 | TCTAGAAAATGTCCTGAAATAAAAAAAATTAGGTGCATTACACTTAAAAA<br>GTTTGTGCAATGCACCTTCGTTAATCAGTATCGGCTCACATTGTCCTGCTT<br>CCCCTGGACATTGCTGTTGGCCCTGTCCTTGCGAGTTCCTTTATTCCAAAG<br>TTCCTCAGAAGCTCCAGGAAGGCGTCTATCTTTTCTGAGTCCCCTGTAACC<br>TCAACTGTCAGGGCATCGGGTGACACATCCACTATCCTCCCCCTGAATATG<br>TTGGTGTACTGTATTATCTCTGACCTTTCTGACTCTGATGGGGCGTGGACC<br>TTGACCATGCAGAGCTCCCTCTTAACTGTTGCTGCGGGTTCAAGGTCCCTG<br>ACCTTTATGACGTCTATGAGCTTGTTCAGCTGCTTTGTTATCTGCTCAAGC<br>ACCCTGTCGTCGCCCCTGGCGATTATTGTCATCCTGGCAATGCCTGGGGTT<br>TCTGATTCCCCGACTGTTATGTTTTCAATGTTGAATCCCCTTCTTGTGAAGA<br>GTCCTGCAACCCTCTGGAAATCGTCTGGTTTGTGCTCCACGAGGGCGCTTA<br>TGATATGGGTATCGGGTTCCATCTCAATCACCATCCGCCTTCCCTGGGGAG<br>TATTTAATTTCCTGGGGGTCCTCCCTTTCAACCCTGTACTCCCCCACTATCT<br>CGGTGAGACCACAGCCGGGGGGGACCATGGGGAGTATCTCATCAGGATCT<br>ATCACTATATCAAGGAGGGCAGGTTCACCTGACCTTATTGCCCTTGAAAG<br>GGCTTCTGAGGTTTCACCGGGTTCCTCTATCCTCTCTGCCTCCACTCCAAA<br>TGATTCTGCCAGCTTCACAAAGTCGGGAACTTCGCCCAGGTGTGTATGGG<br>ACATTCTCTCATCATAGAAGAGCCTCTGCCACTGTGCCACCATTCCAAGGT<br>GCCTGTTGTCCATGATACATATCACCACGGGGATGTCGTATTCCCTTATGG<br>TTGCAAGGTCCTGGCAGACCATGAGGAATCCGCCGTCACCGCACACTGCA<br>ACAACGTCTGAATCAGGCAGTGCCACCTTGGCACCTATGGCGGCTGGAAA<br>ACCGAAGCCCATTGTTCCAAGGCCTCCTGATGATATGAACTTTCTGGGGG<br>CCCTGGATGTGTAGAAGTGGGCCATCCACATCTGGTTCTGTCCCACATCTG<br>TTGTAACGACTGTCTCGTCATCAAGGACCTGGCTTATCTCCTTTATAACCT<br>GCTGGGGCTTCAGGGGCACCTCATCATAGCTCATCCTTGGCATGCAATCG<br>GCCCTGAATTTCTGGACGCTTTCAAGCCACTGGCTGTCCCTCTTTTCATATT<br>TTTTGAGTTTTGCTATGAGTTCCCTGAGGACGTTTCTTGCATCTCCAACGA<br>TGGGGACATCAACCCCAACGTTCTTACCTATCTCTGCGGGGTCGATGTCGA<br>CGTGTATTATCCTGGCGTTGGGGGCGAATTCTGCAACGTTCCCTGTTGTCC<br>TGTCTGAGAATCTGCATCCAACGGCTATGAGGCAGTCGCATTCGTCCACT<br>GTCAGGTTTGCCACCTTCCTGCCGTGCATGCCGAGCATACCCATGGCTGAA<br>GGGTGGTCCTCAGGAAAGGAACCCTTACCAAGGAGTGTTGTTGTCACGGG<br>GGCCTTTATGAGATCTGAGAGTTCCTTTATCTCCCTGGATGCCCCTGATAT<br>TATAACTCCTCCACCTGCAAGTATGACGGGTTTTTCTGACCTCCTTATGAG<br>TTCTGCGGCCCTCTTTATCTGGAGGGGGTGGCCCTTAACATTGGGCCTGTA<br>CCCTGGGAGCTCCAGGTCATCAACCTCCTCCATGATCTCCTGTTCCTGTAT<br>ATCCTTGGGGAGGTCTATAACAACGGGTCCTGGCCTTCCTGTCTTTGCTAT<br>GTGGAAGCTTGCCCTGACAATTGCAGGTATCTCGCTGGCGTCTGATGGCT<br>GGAATGAGTGCTTGGTGATGGGCATGGTTATCCCTATCATGTCCACCTCCT<br>GGAATGCATCATTTCCAATGAGGTGTGTTGGGACCTGACCTGCAATGGCC<br>ACGATGGGGGCTGAGTCCATGTAGGCTGTTGCAATGCCTGTAACAAGGTT<br>TGTTGCCCCGGGACCGGAGGTTGCTATGCAGACCCCCACCCTTCCTGAGG<br>CCCTTGCATATCCGTCTGCTGCGTGTGCTGCGCACTGTTCATGTCTAACGA<br>GGATGTGTTTAAGTTCTGAATCATAGAGCATATCATAGAGTGGCAGGAGC<br>TGTCCACCGGGGTATCCGAAAACGGTGTCTGCTCCCTGATCCAGAAGTGA<br>TCTGATTATTGCCTGGCCACCTTTCATTGGAAACACAATTAACCACCTCAT<br>TTTATGTGATATTATCTATTCATATAATCCTATATAAATATATCGCTAATTT<br>TAAGGTTTTTCTGAGCCATCGGTTGGTTCATGGGGGCGCC | TCTAGAAAATGTCCTGAAATAAAAAAAATTAGGTGCATTACACTTAAAAA<br>GTTTGTGCAATGCACCTTCGTTAATCAGTATCGGCTCACATTGTCCTGCTT<br>CCCCTGGACATTGCTGTTGGCCCTGTCCTTGCGAGTTCCTTTATTCCAAAG<br>TTCCTCAGAAGCTCCAGGAAGGCGTCTATCTTTTCTGAGTCCCCTGTAACC<br>TCAACTGTCAGGGCATCGGGTGACACATCCACTATCCTCCCCCTGAATATG<br>TTGGTGTACTGTATTATCTCTGACCTTTCTGACTCTGATGGGGCGTGGACC<br>TTGACCATGCAGAGCTCCCTCTTAACTGTTGCTGCGGGTTCAAGGTCCCTG<br>ACCTTTATGACGTCTATGAGCTTGTTCAGCTGCTTTGTTATCTGCTCAAGC<br>ACCCTGTCGTCGCCCCTGGCGATTATTGTCATCCTGGCAATGCCTGGGGTT<br>TCTGATTCCCCGACTGTTATGTTTTCAATGTTGAATCCCCTTCTTGTGAAGA<br>GTCCTGCAACCCTCTGGAAATCGTCTGGTTTGTGCTCCACGAGGGCGCTTA<br>TGATATGGGTATCGGGTTCCATCTCAATCACCATCCGCCTTCCCTGGGGAG<br>TATTTAATTTCCTGGGGGTCCTCCCTTTCAACCCTGTACTCCCCCACTATCT<br>CGGTGAGACCACAGCCGGGGGGGACCATGGGGAGTATCTCATCAGGATCT<br>ATCACTATATCAAGGAGGGCAGGTTCACCTGACCTTATTGCCCTTGAAAG<br>GGCTTCTGAGGTTTCACCGGGTTCCTCTATCCTCTCTGCCTCCACTCCAAA<br>TGATTCTGCCAGCTTCACAAAGTCGGGAACTTCGCCCAGGTGTGTATGGG<br>ACATTCTCTCATCATAGAAGAGCCTCTGCCACTGTGCCACCATTCCAAGGT<br>GCCTGTTGTCCATGATACATATCACCACGGGGATGTCGTATTCCCTTATGG<br>TTGCAAGGTCCTGGCAGACCATGAGGAATCCGCCGTCACCGCACACTGCA<br>ACAACGTCTGAATCAGGCAGTGCCACCTTGGCACCTATGGCGGCTGGAAA<br>ACCGAAGCCCATTGTTCCAAGGCCTCCTGATGATATGAACTTTCTGGGGG<br>CCCTGGATGTGTAGAAGTGGGCCATCCACATCTGGTTCTGTCCCACATCTG<br>TTGTAACGACTGTCTCGTCATCAAGGACCTGGCTTATCTCCTTTATAACCT<br>GCTGGGGCTTCAGGGGCACCTCATCATAGCTCATCCTTGGCATGCAATCG<br>GCCCTGAATTTCTGGACGCTTTCAAGCCACTGGCTGTCCCTCTTTTCATATT<br>TTTTGAGTTTTGCTATGAGTTCCCTGAGGACGTTTCTTGCATCTCCAACGA<br>TGGGGACATCAACCCCAACGTTCTTACCTATCTCTGCGGGGTCGATGTCGA<br>CGTGTATTATCCTGGCGTTGGGGGCGAATTCTGCAACGTTCCCTGTTGTCC<br>TGTCTGAGAATCTGCATCCAACGGCTATGAGGCAGTCGCATTCGTCCACT<br>GTCAGGTTTGCCACCTTCCTGCCGTGCATGCCGAGCATACCCATGGCTGAA<br>GGGTGGTCCTCAGGAAAGGAACCCTTACCAAGGAGTGTTGTTGTCACGGG<br>GGCCTTTATGAGATCTGAGAGTTCCTTTATCTCCCTGGATGCCCCTGATAT<br>TATAACTCCTCCACCTGCAAGTATGACGGGTTTTTCTGACCTCCTTATGAG<br>TTCTGCGGCCCTCTTTATCTGGAGGGGGTGGCCCTTAACATTGGGCCTGTA<br>CCCTGGGAGCTCCAGGTCATCAACCTCCTCCATGATCTCCTGTTCCTGTAT<br>ATCCTTGGGGAGGTCTATAACAACGGGTCCTGGCCTTCCTGTCTTTGCTAT<br>GTGGAAGCTTGCCCTGACAATTGCAGGTATCTCGCTGGCGTCTGATGGCT<br>GGAATGAGTGCTTGGTGATGGGCATGGTTATCCCTATCATGTCCACCTCCT<br>GGAATGCATCATTTCCAATGAGGTGTGTTGGGACCTGACCTGCAATGGCC<br>ACGATGGGGGCTGAGTCCATGTAGGCTGTTGCAATGCCTGTAACAAGGTT<br>TGTTGCCCCGGGACCGGAGGTTGCTATGCAGACCCCCACCCTTCCTGAGG<br>CCCTTGCATATCCGTCTGCTGCGTGTGCTGCGCACTGTTCATGTCTAACGA<br>GGATGTGTTTAAGTTCTGAATCATAGAGCATATCATAGAGTGGCAGGAGC<br>TGTCCACCGGGGTATCCGAAAACGGTGTCTGCTCCCTGATCCAGAAGTGA<br>TCTGATTATTGCCTGGCCACCTTTCATTGGAAACACAATTAACCACCTCAT<br>TTTATGTGATATTATCTATTCATATAATCCTATATAAATATATCGCTAATTT<br>TAAGGTTTTTCTGAGCCATCGGTTGGTTCATGGGGGCGCCAGGCTAGGTG<br>GAGGCTCAGTGATGATAAGTCTGCGATGGTGGATGCATGTGTCATGGTCA<br>TAGCTGTTTCCTGTGTGAAATTGTTATCCGCTCAGAGGGCACAATCCTATT<br>CCGCGCTATCCGACAATCTCCAAGACATTAGGTGGAGTTCAGTTCGGCGT<br>ATGGCATATGTCGCTGGAAAGAACATGTGAGCAAAAGGCCAGCAAAAGG<br>CCAGGAACCGTAAAAAGGCCGCGTTGCTGGCGTTTTTCCATAGGCTCCGC<br>CCCCCTGACGAGCATCACAAAAATCGACGCTCAAGTCAGAGGTGGCGAA<br>ACCCGACAGGACTATAAAGATACCAGGCGTTTCCCCCTGGAAGCTCCCTC<br>GTGCGCTCTCCTGTTCCGACCCTGCCGCTTACCGGATACCTGTCCGCCTTT<br>CTCCCTTCGGGAAGCGTGGCGCTTTCTCATAGCTCACGCTGTAGGTATCTC<br>AGTTCGGTGTAGGTCGTTCGCTCCAAGCTGGGCTGTGTGCACGAACCCCC<br>CGTTCAGCCCGACCGCTGCGCCTTATCCGGTAACTATCGTCTTGAGTCCAA<br>CCCGGTAAGACACGACTTATCGCCACTGGCAGCAGCCACTGGTAACAGGA<br>TTAGCAGAGCGAGGTATGTAGGCGGTGCTACAGAGTTCTTGAAGTGGTGG<br>CCTAACTACGGCTACACTAGAAGAACAGTATTTGGTATCTGCGCTCTGCTG<br>AAGCCAGTTACCTTCGGAAAAAGAGTTGGTAGCTCTTGATCCGGCAAACA<br>AACCACCGCTGGTAGCGGTGGTTTTTTTGTTTGCAAGCAGCAGATTACGCG |

| | | | |
|---|---|---|---|
| | | | CAGAAAAAAAGGATCTCAAGAAGATCCTTTGATCTTTTCTACGGGGTCTG<br>ACGCTCTATTCAACAAAGCCGCCGTCCCGTCAAGTCAGCGTAAATGGGTA<br>GGGGGCTTCAAATCGTCCTCGTGATACCAATTCGGAGCCTGCTTTTTTGTA<br>CAAACTTGTTGATAATGGCAATTCAAGGATCTTCACCTAGATCCTTTTAAA<br>TTAAAAATGAAGTTTTAAATCAATCTAAAGTATATATGAGTAAACTTGGT<br>CTGACAGTTACCAATGCTTAATCAGTGAGGCACCTATCTCAGCGATCTGTC<br>TATTTCGTTCATCCATAGTTGCCTGACTCCCCGTCGTGTAGATAACTACGA<br>TACGGGAGGGCTTACCATCTGGCCCCAGTGCTGCAATGATACCGCGAGAG<br>CCACGCTCACCGGCTCCAGATTTATCAGCAATAAACCAGCCAGCCGGAAG<br>GGCCGAGCGCAGAAGTGGTCCTGCAACTTTATCCGCCTCCATCCAGTCTAT<br>TAATTGTTGCCGGGAAGCTAGAGTAAGTAGTTCGCCAGTTAATAGTTTGC<br>GCAACGTTGTTGCCATTGCTACAGGCATCGTGGTGTCACGCTCGTCGTTTG<br>GTATGGCTTCATTCAGCTCCGGTTCCCAACGATCAAGGCGAGTTACATGAT<br>CCCCCATGTTGTGCAAAAAAGCGGTTAGCTCCTTCGGTCCTCCGATCGTTG<br>TCAGAAGTAAGTTGGCCGCAGTGTTATCACTCATGGTTATGGCAGCACTG<br>CATAATTCTCTTACTGTCATGCCATCCGTAAGATGCTTTTCTGTGACTGGT<br>GAGTACTCAACCAAGTCATTCTGAGAATAGTGTATGCGGCGACCGAGTTG<br>CTCTTGCCCGGCGTCAATACGGGATAATACCGCGCCACATAGCAGAACTT<br>TAAAAGTGCTCATCATTGGAAAACGTTCTTCGGGGCGAAAACTCTCAAGG<br>ATCTTACCGCTGTTGAGATCCAGTTCGATGTAACCCACTCGTGCACCCAAC<br>TGATCTTCAGCATCTTTTACTTTCACCAGCGTTTCTGGGTGAGCAAAAACA<br>GGAAGGCAAAATGCCGCAAAAAAGGGAATAAGGGCGACACGGAAATGTT<br>GAATACTCATACTCTTCCTTTTTCAATATTATTGAAGCATTTATCAGGGTT<br>ATTGTCTCATGAGCGGATACATATTTGAATGTATTTAGAAAAATAAACAA<br>ATAGGGGTTCCGCGCACATTTCCCCGAAAAGTGCCAGATACCTGAAACAA<br>AACCCATCGTACGGCCAAGGAAGTCTCCAATAACTGTGATCCACCACAAG<br>CGCCAGGGTTTTCCCAGTCACGACGTTGTAAAACGACGGCCAGTCATGCA<br>TAATCCGCACGCATCTGGAATAAGGAAGTGCCATTCCGCCTGACCT |
| Gene | TetR codon optimized | CACTTAAACGGCCGGCCCTACGGATTCCCCATGAACCAACCGATGGCTCA<br>GAAAAACCTTAAAATTAGCGATATATTTATATAGGATTATATGAATAGAT<br>AATATCACATAAAATGAGGTGGTTAATTATGTCCAGACTCGATAAATCAA<br>AAGTGATTAACAGCGCACTCGAGCTGCTTAATGAGGTCGGAATCGAAGGT<br>CTCACAACCAGGAAACTCGCCCAGAAGCTCGGTGTTGAGCAGCCTACACT<br>CTATTGGCATGTTAAAAATAAGAGGGCACTCCTCGACGCCCTCGCCATTG<br>AGATGCTCGATAGGCACCATACACACTTTTGCCCTCTCGAAGGGGAAAGC<br>TGGCAGGATTTTCTCAGGAATAACGCAAAATCATTTAGATGTGCACTCCTC<br>TCACATAGGGATGGAGCAAAAGTTCATCTCGGTACAAGGCCTACAGAAAA<br>ACAGTATGAAACACTCGAAAATCAGCTCGCCTTTCTCTGCCAGCAGGGTT<br>TTTCACTCGAGAATGCACTCTATGCACTCAGCGCAGTGGGGCATTTTACAC<br>TCGGTTGCGTTCTCGAAGATCAGGAGCATCAGGTCGCAAAAGAAGAAAG<br>GGAAACACCTACAACAGATTCAATGCCCCCACTCCTCAGGCAGGCAATCG<br>AACTCTTTGATCACCAGGGTGCAGAGCCAGCCTTCCTCTTCGGCCTTGAAC<br>TCATCATATGCGGACTCGAAAAACAGCTTAAATGTGAATCAGGGTCCTAA<br>CTATTGCCAGGCGATCGCTGATTCACTG | CACTTAAACGGCCGGCCCTACGGATTCCCCATGAACCAACCGATGGCTCA<br>GAAAAACCTTAAAATTAGCGATATATTTATATAGGATTATATGAATAGAT<br>AATATCACATAAAATGAGGTGGTTAATTATGTCCAGACTCGATAAATCAA<br>AAGTGATTAACAGCGCACTCGAGCTGCTTAATGAGGTCGGAATCGAAGGT<br>CTCACAACCAGGAAACTCGCCCAGAAGCTCGGTGTTGAGCAGCCTACACT<br>CTATTGGCATGTTAAAAATAAGAGGGCACTCCTCGACGCCCTCGCCATTG<br>AGATGCTCGATAGGCACCATACACACTTTTGCCCTCTCGAAGGGGAAAGC<br>TGGCAGGATTTTCTCAGGAATAACGCAAAATCATTTAGATGTGCACTCCTC<br>TCACATAGGGATGGAGCAAAAGTTCATCTCGGTACAAGGCCTACAGAAAA<br>ACAGTATGAAACACTCGAAAATCAGCTCGCCTTTCTCTGCCAGCAGGGTT<br>TTTCACTCGAGAATGCACTCTATGCACTCAGCGCAGTGGGGCATTTTACAC<br>TCGGTTGCGTTCTCGAAGATCAGGAGCATCAGGTCGCAAAAGAAGAAAG<br>GGAAACACCTACAACAGATTCAATGCCCCCACTCCTCAGGCAGGCAATCG<br>AACTCTTTGATCACCAGGGTGCAGAGCCAGCCTTCCTCTTCGGCCTTGAAC<br>TCATCATATGCGGACTCGAAAAACAGCTTAAATGTGAATCAGGGTCCTAA<br>CTATTGCCAGGCGATCGCTGATTCACTGAGGCTAGGTGGAGGCTCAGTGA<br>TGATAAGTCTGCGATGGTGGATGCATGTGTCATGGTCATAGCTGTTTCCTG<br>TGTGAAATTGTTATCCGCTCAGAGGGCACAATCCTATTCCGCGCTATCCGA<br>CAATCTCCAAGACATTAGGTGGAGTTCAGTTCGGCGTATGGCATATGTCG<br>CTGGAAAGAACATGTGAGCAAAAGGCCAGCAAAAGGCCAGGAACCGTAA<br>AAAGGCCGCGTTGCTGGCGTTTTTCCATAGGCTCCGCCCCCCTGACGAGC<br>ATCACAAAAATCGACGCTCAAGTCAGAGGTGGCGAAACCCGACAGGACT<br>ATAAAGATACCAGGCGTTTCCCCCTGGAAGCTCCCTCGTGCGCTCTCCTGT<br>TCCGACCCTGCCGCTTACCGGATACCTGTCCGCCTTTCTCCCTTCGGGAAG<br>CGTGGCGCTTTCTCATAGCTCACGCTGTAGGTATCTCAGTTCGGTGTAGGT<br>CGTTCGCTCCAAGCTGGGCTGTGTGCACGAACCCCCCGTTCAGCCCGACC<br>GCTGCGCCTTATCCGGTAACTATCGTCTTGAGTCCAACCCGGTAAGACAC<br>GACTTATCGCCACTGGCAGCAGCCACTGGTAACAGGATTAGCAGAGCGAG<br>GTATGTAGGCGGTGCTACAGAGTTCTTGAAGTGGTGGCCTAACTACGGCT<br>ACACTAGAAGAACAGTATTTGGTATCTGCGCTCTGCTGAAGCCAGTTACC<br>TTCGGAAAAAGAGTTGGTAGCTCTTGATCCGGCAAACAAACCACCGCTGG<br>TAGCGGTGGTTTTTTTGTTTGCAAGCAGCAGATTACGCGCAGAAAAAAAG<br>GATCTCAAGAAGATCCTTTGATCTTTTCTACGGGGTCTGACGCTCTATTCA<br>ACAAAGCCGCCGTCCCGTCAAGTCAGCGTAAATGGGTAGGGGGCTTCAAA<br>TCGTCCTCGTGATACCAATTCGGAGCCTGCTTTTTTGTACAAACTTGTTGA<br>TAATGGCAATTCAAGGATCTTCACCTAGATCCTTTTAAATTAAAAATGAA<br>GTTTTAAATCAATCTAAAGTATATATGAGTAAACTTGGTCTGACAGTTACC<br>AATGCTTAATCAGTGAGGCACCTATCTCAGCGATCTGTCTATTTCGTTCAT<br>CCATAGTTGCCTGACTCCCCGTCGTGTAGATAACTACGATACGGGAGGGC |

| | | | |
|---|---|---|---|
| | | | TTACCATCTGGCCCCAGTGCTGCAATGATACCGCGAGAGCCACGCTCACC<br>GGCTCCAGATTTATCAGCAATAAACCAGCCAGCCGGAAGGGCCGAGCGC<br>AGAAGTGGTCCTGCAACTTTATCCGCCTCCATCCAGTCTATTAATTGTTGC<br>CGGGAAGCTAGAGTAAGTAGTTCGCCAGTTAATAGTTTGCGCAACGTTGT<br>TGCCATTGCTACAGGCATCGTGGTGTCACGCTCGTCGTTTGGTATGGCTTC<br>ATTCAGCTCCGGTTCCCAACGATCAAGGCGAGTTACATGATCCCCCATGTT<br>GTGCAAAAAAGCGGTTAGCTCCTTCGGTCCTCCGATCGTTGTCAGAAGTA<br>AGTTGGCCGCAGTGTTATCACTCATGGTTATGGCAGCACTGCATAATTCTC<br>TTACTGTCATGCCATCCGTAAGATGCTTTTCTGTGACTGGTGAGTACTCAA<br>CCAAGTCATTCTGAGAATAGTGTATGCGGCGACCGAGTTGCTCTTGCCCG<br>GCGTCAATACGGGATAATACCGCGCCACATAGCAGAACTTTAAAAGTGCT<br>CATCATTGGAAAACGTTCTTCGGGGCGAAAACTCTCAAGGATCTTACCGC<br>TGTTGAGATCCAGTTCGATGTAACCCACTCGTGCACCCAACTGATCTTCAG<br>CATCTTTTACTTTCACCAGCGTTTCTGGGTGAGCAAAAACAGGAAGGCAA<br>AATGCCGCAAAAAAGGGAATAAGGGCGACACGGAAATGTTGAATACTCA<br>TACTCTTCCTTTTTCAATATTATTGAAGCATTTATCAGGGTTATTGTCTCAT<br>GAGCGGATACATATTTGAATGTATTTAGAAAAATAAACAAATAGGGGTTC<br>CGCGCACATTTCCCCGAAAAGTGCCAGATACCTGAAACAAAACCCATCGT<br>ACGGCCAAGGAAGTCTCCAATAACTGTGATCCACCACAAGCGCCAGGGTT<br>TTCCCAGTCACGACGTTGTAAAACGACGGCCAGTCATGCATAATCCGCAC<br>GCATCTGGAATAAGGAAGTGCCATTCCGCCTGACCT |
| Promoter | SIP1 | GGCGCCTAAATCACCGTGAATTTGCGAATAATTTCATAATTATCTGAAAA<br>GTATATCAAACTATCCAAATATTAAAAATTACCGCCATTTAAAATATATTG<br>AATGTACCGTAAACTATATATATTGAAAAAGCGGAAAGCTATTCCCTATC<br>AGTGATAGAGAGGAGGAATATATGTGTTTCCAATGAAAGGTGGCCAGGC<br>AATAATCAGATCACTTCTGGATCAGGGAGCAGACACCGTTTTCGGATACC<br>CCGGTGGACAGCTCCTGCCACTCTATGATATGCTCTATGATTCAGAACTTA<br>AACACATCCTCGTTAGACATGAACAGTGCGCAGCACACGCAGCAGACGG<br>ATATGCAAGGGCCTCAGGAAGGGTGGGGGTCTGCATAGCAACCTCCGGTC<br>CCGGTGCAACAAACCTTGTTACAGGCATTGCAACAGCCTACATGGACTCA<br>GCCCCCATCGTGGCCATTGCAGGTCAGGTCCCAACACACCTCATTGGAAA<br>TGATGCATTCCAGGAGGTGGACATGATAGGGATAACCATGCCCATCACCA<br>AGCACTCATTCCAGCCATCAGACGCCAGCGAGATACCTGCAATTGTC | CAATCCGCCCTCACTACAACCGGGCGCCTAAATCACCGTGAATTTGCGAA<br>TAATTTCATAATTATCTGAAAAGTATATCAAACTATCCAAATATTAAAAAT<br>TACCGCCATTTAAAATATATTGAATGTACCGTAAACTATATATATTGAAAA<br>AGCGGAAAGCTATTCCCTATCAGTGATAGAGAGGAGGAATATATGTGTTT<br>CCAATGAAAGGTGGCCAGGCAATAATCAGATCACTTCTGGATCAGGGAGC<br>AGACACCGTTTTCGGATACCCCGGTGGACAGCTCCTGCCACTCTATGATAT<br>GCTCTATGATTCAGAACTTAAACACATCCTCGTTAGACATGAACAGTGCG<br>CAGCACACGCAGCAGACGGATATGCAAGGGCCTCAGGAAGGGTGGGGGT<br>CTGCATAGCAACCTCCGGTCCCGGTGCAACAAACCTTGTTACAGGCATTG<br>CAACAGCCTACATGGACTCAGCCCCCATCGTGGCCATTGCAGGTCAGGTC<br>CCAACACACCTCATTGGAAATGATGCATTCCAGGAGGTGGACATGATAGG<br>GATAACCATGCCCATCACCAAGCACTCATTCCAGCCATCAGACGCCAGCG<br>AGATACCTGCAATTGTCCTACTCTGGCGTCGATGAGGGA |
| Promoter | SIP2 | GGCGCCTAAATCACCGTGAATTTGCGAATAATTTCATAATTATCTGAAAA<br>GTATATCAAACTATCCAAATATTAAAAATTACCGCCATTTAAAATATATTG<br>AATGTACCGTAAACTATATATATTGAAAAAGCCCCTATCAGTGATAGAGA<br>AAGATGATTTTGGAGGAATATATGTGTTTCCAATGAAAGGTGGCCAGGCA<br>ATAATCAGATCACTTCTGGATCAGGGAGCAGACACCGTTTTCGGATACCC<br>CGGTGGACAGCTCCTGCCACTCTATGATATGCTCTATGATTCAGAACTTAA<br>ACACATCCTCGTTAGACATGAACAGTGCGCAGCACACGCAGCAGACGGAT<br>ATGCAAGGGCCTCAGGAAGGGTGGGGGTCTGCATAGCAACCTCCGGTCCC<br>GGTGCAACAAACCTTGTTACAGGCATTGCAACAGCCTACATGGACTCAGC<br>CCCCATCGTGGCCATTGCAGGTCAGGTCCCAACACACCTCATTGGAAATG<br>ATGCATTCCAGGAGGTGGACATGATAGGGATAACCATGCCCATCACCAAG<br>CACTCATTCCAGCCATCAGACGCCAGCGAGATACCTGCAATTGTC | CAATCCGCCCTCACTACAACCGGGCGCCTAAATCACCGTGAATTTGCGAA<br>TAATTTCATAATTATCTGAAAAGTATATCAAACTATCCAAATATTAAAAAT<br>TACCGCCATTTAAAATATATTGAATGTACCGTAAACTATATATATTGAAAA<br>AGCCCCTATCAGTGATAGAGAAAGATGATTTTGGAGGAATATATGTGTTT<br>CCAATGAAAGGTGGCCAGGCAATAATCAGATCACTTCTGGATCAGGGAGC<br>AGACACCGTTTTCGGATACCCCGGTGGACAGCTCCTGCCACTCTATGATAT<br>GCTCTATGATTCAGAACTTAAACACATCCTCGTTAGACATGAACAGTGCG<br>CAGCACACGCAGCAGACGGATATGCAAGGGCCTCAGGAAGGGTGGGGGT<br>CTGCATAGCAACCTCCGGTCCCGGTGCAACAAACCTTGTTACAGGCATTG<br>CAACAGCCTACATGGACTCAGCCCCCATCGTGGCCATTGCAGGTCAGGTC<br>CCAACACACCTCATTGGAAATGATGCATTCCAGGAGGTGGACATGATAGG<br>GATAACCATGCCCATCACCAAGCACTCATTCCAGCCATCAGACGCCAGCG<br>AGATACCTGCAATTGTCCTACTCTGGCGTCGATGAGGGA |
| Promot | TATA | GGCGCCCCCATGAACCAACCGATGGCTCAGAAAAACCTTAAAATTAGCGA<br>TATATTTATATAGGCCCTATCAGTGATAGAGAATCACATAAAATGAGGTG<br>GTTAATTGTGTTTCCAATGAAAGGTGGCCAGGCAATAATCAGATCACTTCT<br>GGATCAGGGAGCAGACACCGTTTTCGGATACCCCGGTGGACAGCTCCTGC<br>CACTCTATGATATGCTCTATGATTCAGAACTTAAACACATCCTCGTTAGAC<br>ATGAACAGTGCGCAGCACACGCAGCAGACGGATATGCAAGGGCCTCAGG<br>AAGGGTGGGGGTCTGCATAGCAACCTCCGGTCCCGGTGCAACAAACCTTG<br>TTACAGGCATTGCAACAGCCTACATGGACTCAGCCCCCATCGTGGCCATT<br>GCAGGTCAGGTCCCAACACACCTCATTGGAAATGATGCATTCCAGGAGGT<br>GGACATGATAGGGATAACCATGCCCATCACCAAGCACTCATTCCAGCCAT<br>CAGACGCCAGCGAGATACCTGCAATTGTC | CAATCCGCCCTCACTACAACCGGGCGCCCCCATGAACCAACCGATGGCTC<br>AGAAAAACCTTAAAATTAGCGATATATTTATATAGGCCCTATCAGTGATA<br>GAGAATCACATAAAATGAGGTGGTTAATTGTGTTTCCAATGAAAGGTGGC<br>CAGGCAATAATCAGATCACTTCTGGATCAGGGAGCAGACACCGTTTTCGG<br>ATACCCCGGTGGACAGCTCCTGCCACTCTATGATATGCTCTATGATTCAGA<br>ACTTAAACACATCCTCGTTAGACATGAACAGTGCGCAGCACACGCAGCAG<br>ACGGATATGCAAGGGCCTCAGGAAGGGTGGGGGTCTGCATAGCAACCTCC<br>GGTCCCGGTGCAACAAACCTTGTTACAGGCATTGCAACAGCCTACATGGA<br>CTCAGCCCCCATCGTGGCCATTGCAGGTCAGGTCCCAACACACCTCATTG<br>GAAATGATGCATTCCAGGAGGTGGACATGATAGGGATAACCATGCCCATC<br>ACCAAGCACTCATTCCAGCCATCAGACGCCAGCGAGATACCTGCAATTGT<br>CCTACTCTGGCGTCGATGAGGGA |

| | | | |
|---|---|---|---|
| er | | | |
| Promoter | Trans | GGCGCCCCCATGAACCAACCGATGGCTCAGAAAAACCTTAAAATTAGCGA<br>TATATTTATATAGGATTATATGAATAGATAATATCACACCCTATCAGTGAT<br>AGAGAAGGTGGTTAATTGTGTTTCCAATGAAAGGTGGCCAGGCAATAATC<br>AGATCACTTCTGGATCAGGGAGCAGACACCGTTTTCGGATACCCCGGTGG<br>ACAGCTCCTGCCACTCTATGATATGCTCTATGATTCAGAACTTAAACACAT<br>CCTCGTTAGACATGAACAGTGCGCAGCACACGCAGCAGACGGATATGCAA<br>GGGCCTCAGGAAGGGTGGGGGTCTGCATAGCAACCTCCGGTCCCGGTGCA<br>ACAAACCTTGTTACAGGCATTGCAACAGCCTACATGGACTCAGCCCCCAT<br>CGTGGCCATTGCAGGTCAGGTCCCAACACACCTCATTGGAAATGATGCAT<br>TCCAGGAGGTGGACATGATAGGGATAACCATGCCCATCACCAAGCACTCA<br>TTCCAGCCATCAGACGCCAGCGAGATACCTGCAATTGTC | CAATCCGCCCTCACTACAACCGGGCGCCCCCATGAACCAACCGATGGCTC<br>AGAAAAACCTTAAAATTAGCGATATATTTATATAGGATTATATGAATAGA<br>TAATATCACACCCTATCAGTGATAGAGAAGGTGGTTAATTGTGTTTCCAAT<br>GAAAGGTGGCCAGGCAATAATCAGATCACTTCTGGATCAGGGAGCAGAC<br>ACCGTTTTCGGATACCCCGGTGGACAGCTCCTGCCACTCTATGATATGCTC<br>TATGATTCAGAACTTAAACACATCCTCGTTAGACATGAACAGTGCGCAGC<br>ACACGCAGCAGACGGATATGCAAGGGCCTCAGGAAGGGTGGGGGTCTGC<br>ATAGCAACCTCCGGTCCCGGTGCAACAAACCTTGTTACAGGCATTGCAAC<br>AGCCTACATGGACTCAGCCCCCATCGTGGCCATTGCAGGTCAGGTCCCAA<br>CACACCTCATTGGAAATGATGCATTCCAGGAGGTGGACATGATAGGGATA<br>ACCATGCCCATCACCAAGCACTCATTCCAGCCATCAGACGCCAGCGAGAT<br>ACCTGCAATTGTCCTACTCTGGCGTCGATGAGGGA |
| Promoter | TATA_Trans | GGCGCCCCCATGAACCAACCGATGGCTCAGAAAAACCTTAAAATTAGCGA<br>TATATTTATATAGGCCCTATCAGTGATAGAGAATCACACCCTATCAGTGAT<br>AGAGAAGGTGGTTAATTGTGTTTCCAATGAAAGGTGGCCAGGCAATAATC<br>AGATCACTTCTGGATCAGGGAGCAGACACCGTTTTCGGATACCCCGGTGG<br>ACAGCTCCTGCCACTCTATGATATGCTCTATGATTCAGAACTTAAACACAT<br>CCTCGTTAGACATGAACAGTGCGCAGCACACGCAGCAGACGGATATGCAA<br>GGGCCTCAGGAAGGGTGGGGGTCTGCATAGCAACCTCCGGTCCCGGTGCA<br>ACAAACCTTGTTACAGGCATTGCAACAGCCTACATGGACTCAGCCCCCAT<br>CGTGGCCATTGCAGGTCAGGTCCCAACACACCTCATTGGAAATGATGCAT<br>TCCAGGAGGTGGACATGATAGGGATAACCATGCCCATCACCAAGCACTCA<br>TTCCAGCCATCAGACGCCAGCGAGATACCTGCAATTGTC | CAATCCGCCCTCACTACAACCGGGCGCCCCCATGAACCAACCGATGGCTC<br>AGAAAAACCTTAAAATTAGCGATATATTTATATAGGCCCTATCAGTGATA<br>GAGAATCACACCCTATCAGTGATAGAGAAGGTGGTTAATTGTGTTTCCAA<br>TGAAAGGTGGCCAGGCAATAATCAGATCACTTCTGGATCAGGGAGCAGAC<br>ACCGTTTTCGGATACCCCGGTGGACAGCTCCTGCCACTCTATGATATGCTC<br>TATGATTCAGAACTTAAACACATCCTCGTTAGACATGAACAGTGCGCAGC<br>ACACGCAGCAGACGGATATGCAAGGGCCTCAGGAAGGGTGGGGGTCTGC<br>ATAGCAACCTCCGGTCCCGGTGCAACAAACCTTGTTACAGGCATTGCAAC<br>AGCCTACATGGACTCAGCCCCCATCGTGGCCATTGCAGGTCAGGTCCCAA<br>CACACCTCATTGGAAATGATGCATTCCAGGAGGTGGACATGATAGGGATA<br>ACCATGCCCATCACCAAGCACTCATTCCAGCCATCAGACGCCAGCGAGAT<br>ACCTGCAATTGTCCTACTCTGGCGTCGATGAGGGA |

36

### Supplementary Figures

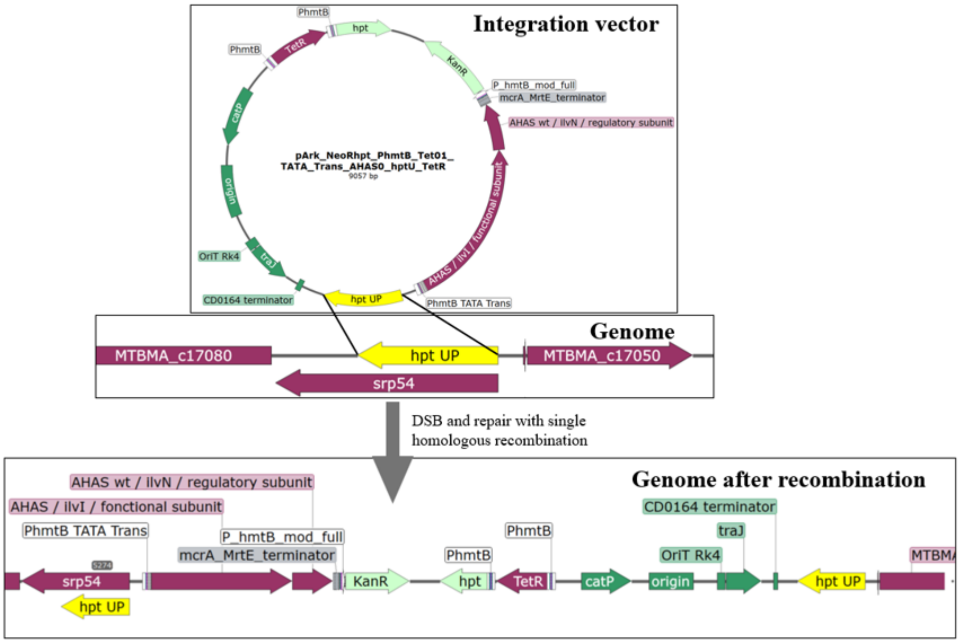


**Supplementary figure S1**. Example of vector construct and genome integration for the TATA_Trans, a $P_{hmtB}$ based promoter. The elements for *E. coli* replication, selection and mobilization are shown in darker green; selection marker for *M. marburgensis* is shown in light green. The flank for homologous recombination upstream of the hypoxanthine phosphoribosyltransferase gene (*hpt*) is shown in yellow. Functional genes or genes of interest in *M. marburgensis* are shown in purple. DSB stands for double strand break and happens spontaneously.

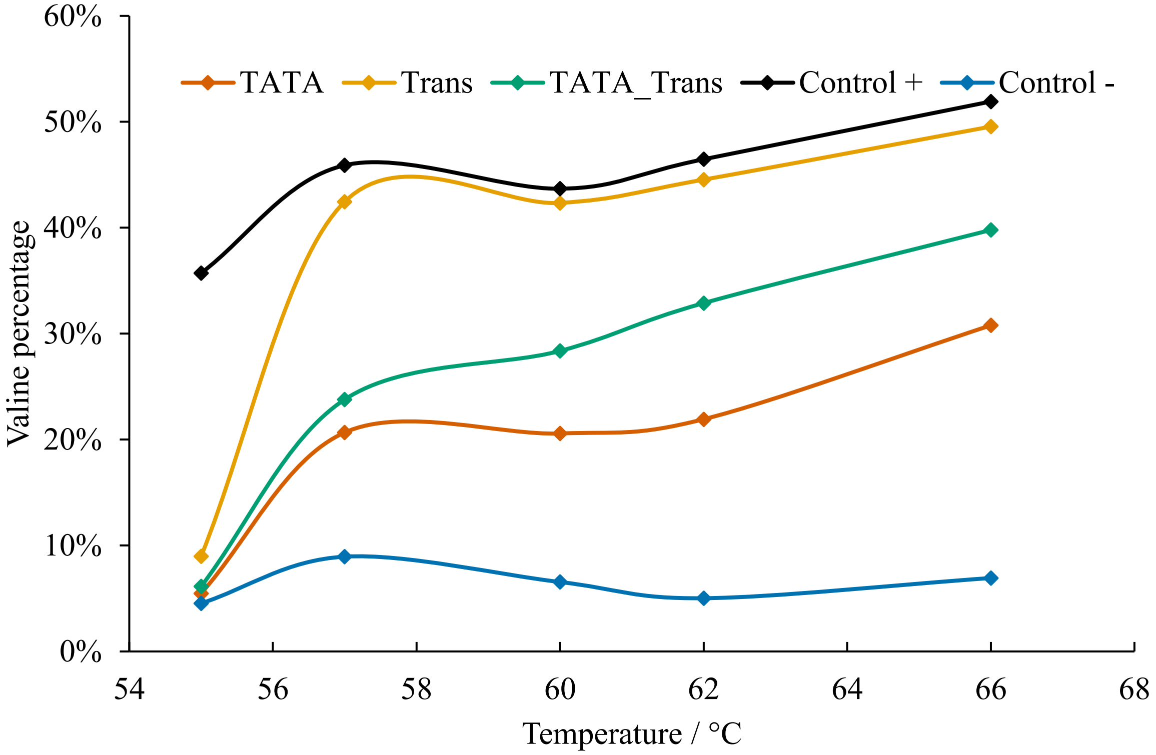


**Supplementary figure S2**. Ratio of valine compared to all 20 proteinogenic amino acids measured in the supernatant for each *M. marburgensis* strain containing a promoter (TATA, TATA_Trans, Trans, $Ctrl^+$, $Ctrl^-$) expressing the AHAS gene after overnight incubation. $Control^+$ is the native, constitutively expressed AHAS with $P_{hmtB}$ and $Control^-$ is the *M. marburgensis* strain with no promoter in front of AHAS (n=1).

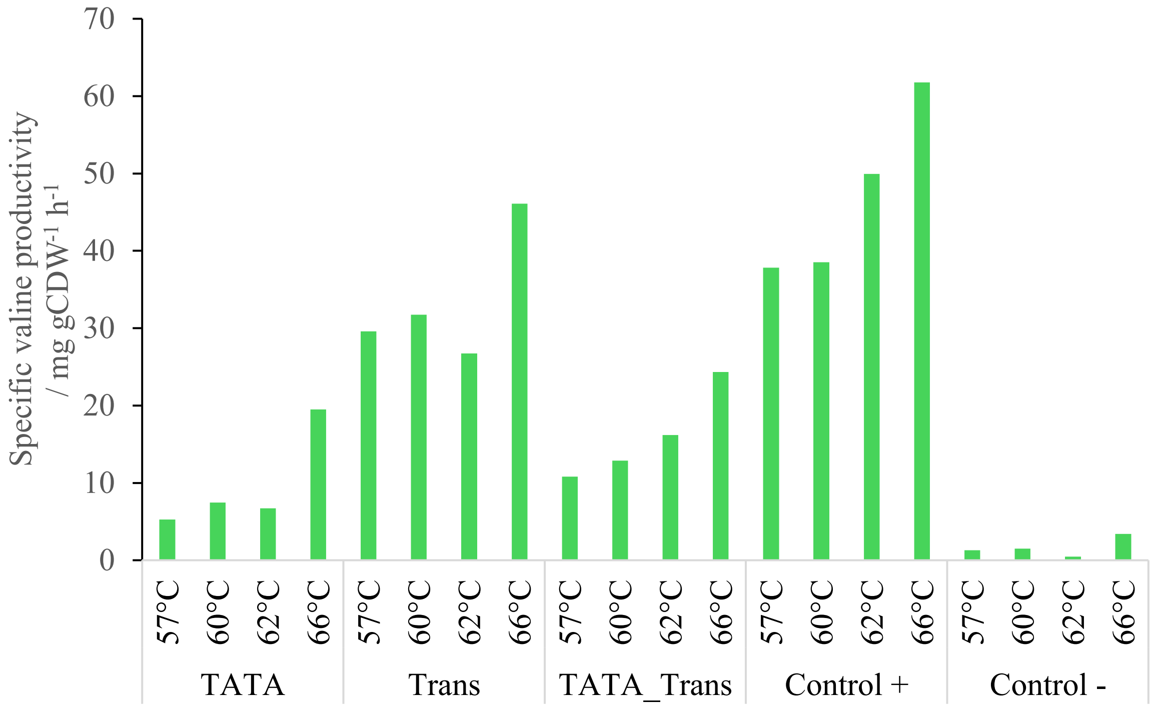


**Supplementary figure S3**. Specific valine productivities for each *M. marburgensis* strain with promoter (TATA, TATA_Trans, Trans, Ctrl$^{+}$, Ctrl$^{-}$) expressing the native AHAS encoding gene of *Methanothermobacter thermautotrophicus*. Control $^{+}$ is the native, constitutively expressed, $P_{hmtB}$ and Control $^{-}$ is the *M. marburgensis* strain with no promoter in front of the AHAS encoding (n=1).

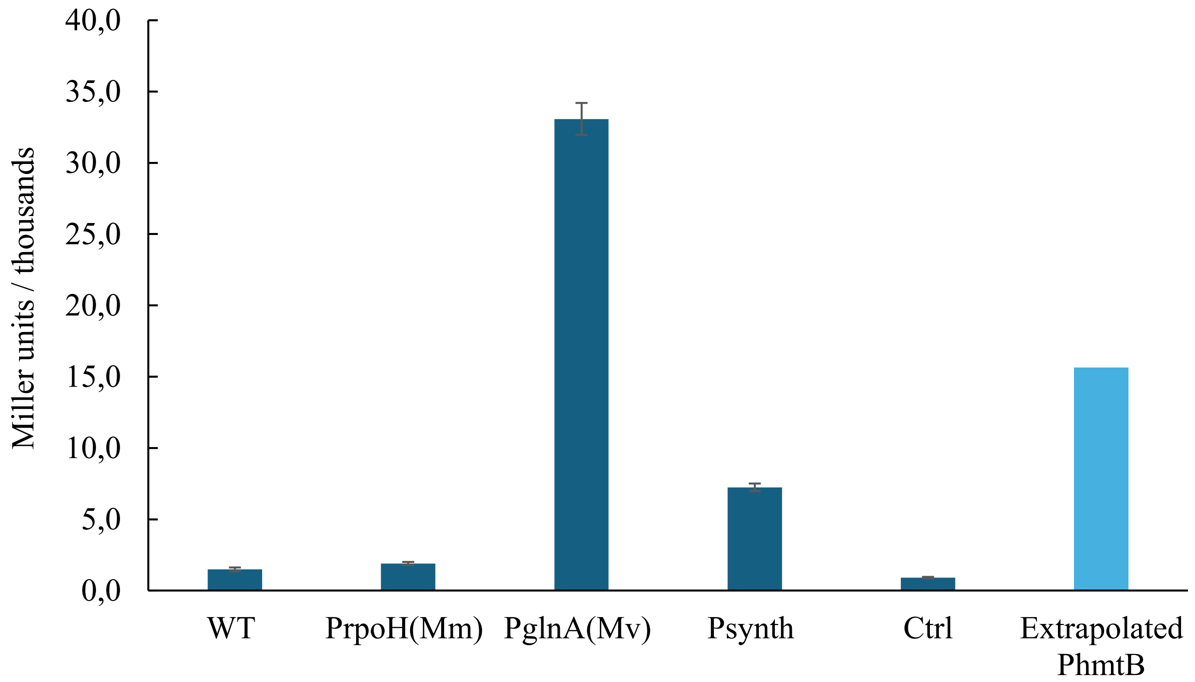


**Supplementary figure S4**. Enzyme activity of the thermostable betagalactosidase (Bgab) in Miller units produced with *M. marburgensis* containing various promoters ($P_{rpoH}$ from *M. marburgensis* (PrpoH(Mm)), $P_{glnA}$ from *Methanococcus vanielli* ($P_{glnA(Mv)}$), $P_{synth}$, and extrapolated $P_{hmtB}$ (Fink et al. 2021)) initiating *bgab* expression. Additional controls were performed, such as sterile *M. marburgensis*

medium (Ctrl) and crude extract from wild-type *M. marburgensis* (WT). Activity was tested with ONPG assay at 420 nm and shown as Miller Units (n=6). Error bars indicate standard deviation.

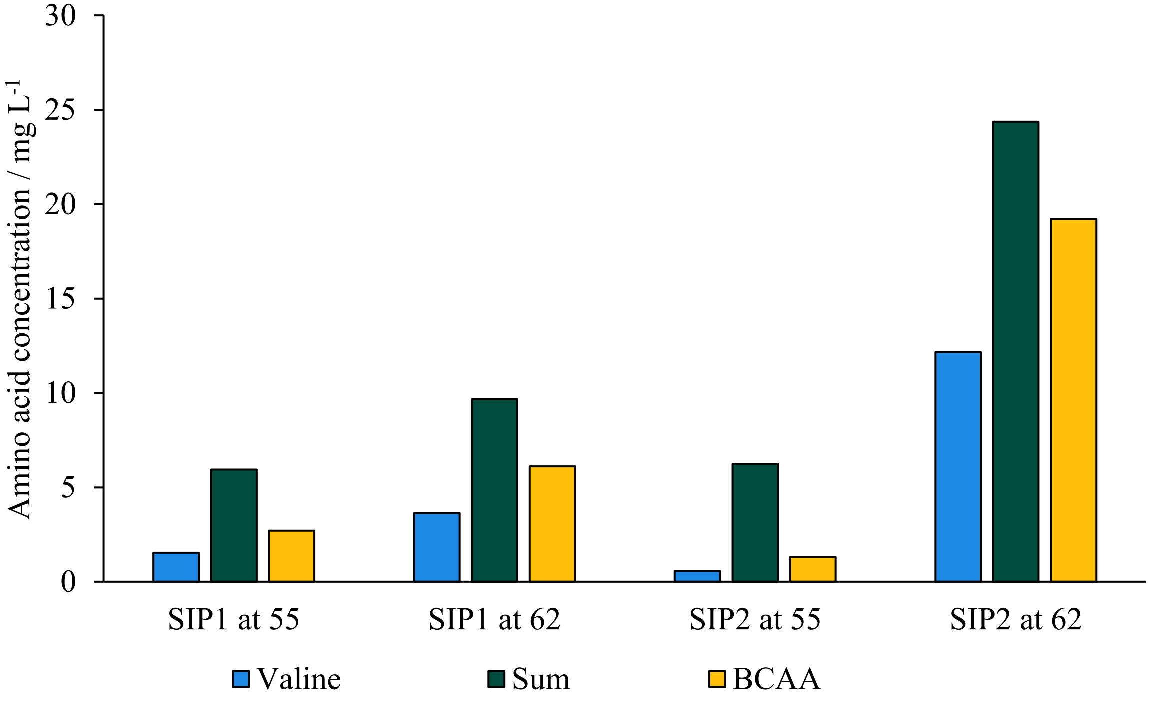


**Supplementary figure S5**. Concentration of different single amino acids and mixtures thereof measured in the supernatant of overnight cultures of *M. marburgensis* strains with two engineered promoters (SIP1, SIP2) expressing the native AHAS encoding gene from *M. thermautotrophicus* (n=1). BCAA stands for branched-chain amino acids. Numbers indicate growth temperature in degree Celsius.

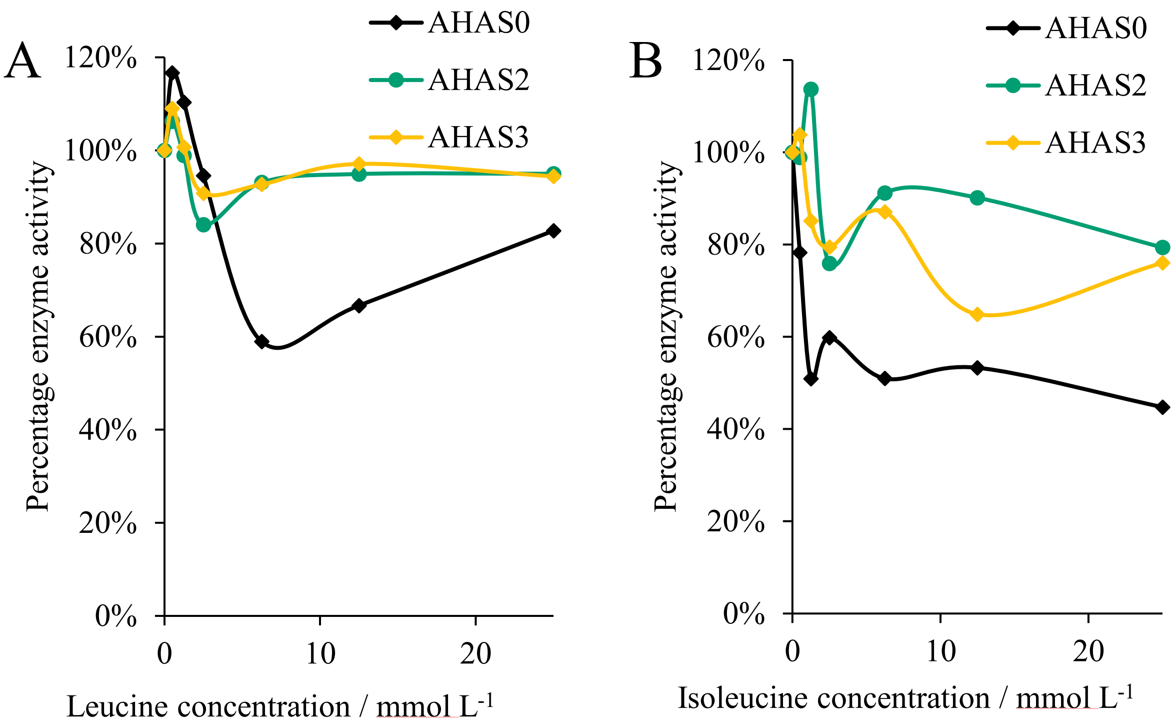


**Supplementary figure S6**. Activity of each AHAS variant tested in *M. marburgensis* initiated with MIP analyzed using different leucine and isoleucine concentrations as putative allosteric inhibitors. Results shown relative to the AHAS activity without leucine or valine (n=2)

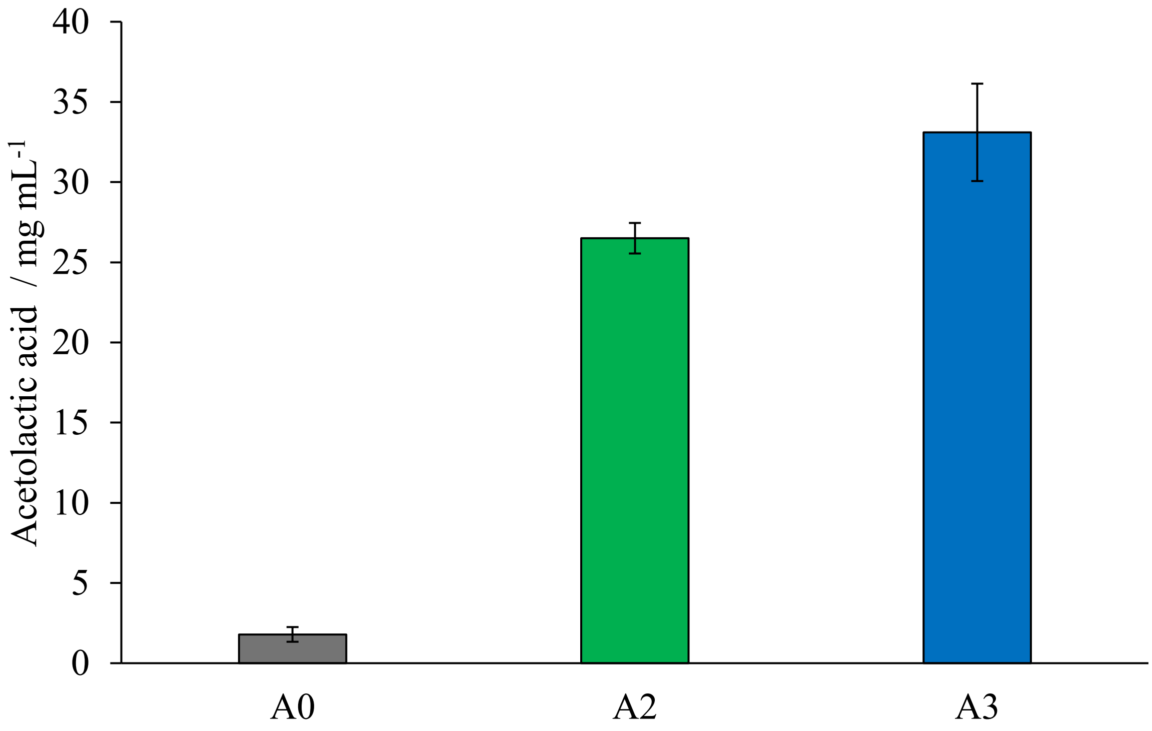


**Supplementary figure S7**. Acetolactic acid concentration in supernatant after overnight growth at 62 °C (ON state). All genes are under the MIP promoter. A0: AHAS0, wild type enzyme.; A2: AHAS2; A3: AHAS3. Standard deviation given as error bars (n=2)

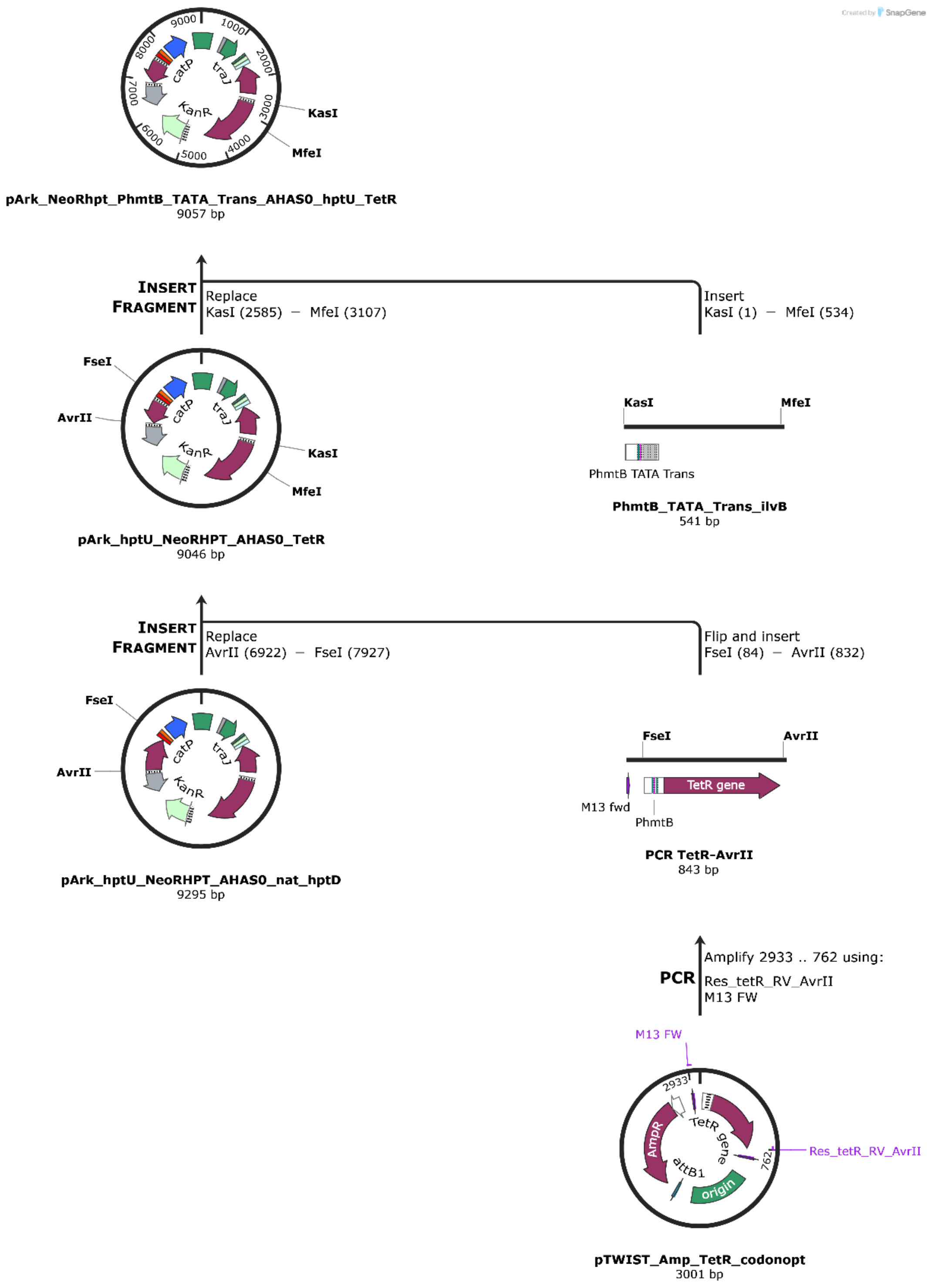


**Supplementary figure S8**. Example of steps performed to obtain the vector constructs used for transformation of *M. marburgensis*. Depending on the final construct, the starting plasmid might contain AHAS0, AHAS2 or AHAS3 and the fragment used to replace the promoter might be $P_{hmtB}$_TATA_Trans_ilvB, $P_{hmtB}$_TATA_ilvB, $P_{hmtB}$_Trans_ilvB, SIP1_ilvB or SIP2_ilvB. The part of AHAS added with the promoter had one codon change to remove *Sma*I restriction enzyme recognition site to allow for quick recognition of the right constructs with *Sma*I restriction digest.
